# Attention-like regulation of theta sweeps in the brain’s spatial navigation circuit

**DOI:** 10.64898/2026.01.27.702083

**Authors:** Abraham Z. Vollan, Michael F. Schellenberger, May-Britt Moser, Edvard I. Moser

**Author notes:** Corresponding authors:,; Editorial correspondence.

## Abstract

Spatial attention supports navigation by prioritizing information from selected locations. A candidate neural mechanism is provided by theta-paced sweeps in grid– and place-cell population activity, which sample nearby space in a left-right-alternating pattern coordinated by parasubicular direction signals. During exploration, this alternation promotes uniform spatial coverage, but whether sweeps can be flexibly tuned to locations of particular interest remains unclear. Using large-scale Neuropixels recordings in freely-behaving rats, we show that sweeps and direction signals are rapidly and dynamically modulated: they track moving targets during pursuit, precede orienting responses during immobility, and reverse during backward locomotion—without prior spatial learning. Similar modulation occurs during REM sleep. Canonical head-direction signals remain head-aligned. These findings identify sweeps as a flexible, attention-like mechanism for selectively sampling allocentric cognitive maps.

## Main Text

Neurons in the hippocampus and medial entorhinal cortex (MEC) form internal representations of space that support navigation and memory (1–5). Beyond tracking the animal’s instantaneous position, population activity in this circuit is organized within individual cycles of the theta rhythm to sample time-compressed sequences of locations ahead of the animal (6–11). These theta-paced sequences, or sweeps, emanate from the animal’s current location and alternate in a stereotyped left-right pattern across successive theta cycles (11). Sweep direction is coordinated by input from internal direction cells in parasubiculum (11), which differ from canonical head direction cells in other areas (12, 13) in that they exhibit theta-cycle skipping (14–16), are more broadly tuned to head direction (17, 18), and, at the population level, encode an ‘internal direction signal’ that alternates left and right relative to the animal’s heading with offsets of ∼30° to either side (11). When imposed on grid– and place-cell sweeps, this directional alternation generates a broad, forward-oriented sampling sector that maximizes uniform spatial coverage and may promote efficient map formation (11, 19–21).

Natural navigation, however, often imposes demands that are poorly served by uniform sampling, such as tracking moving targets, recovering orientation after target loss, planning routes around obstacles, or navigating to learned goals. It remains unclear whether entorhinal-hippocampal sweeps can be modulated to meet such demands. Existing data leave unresolved whether sweeps are inherently constrained to fixed left-right angles (11) or to learned trajectories to familiar targets (9, 10, 22–27), or whether they can instead be flexibly steered in any direction to support the unpredictable requirements of real-world navigation, in line with grid cell-based computational models (28–30). Here, we propose that entorhinal–hippocampal sweeps implement a form of internal spatial attention: a mechanism that dynamically allocates sampling toward any location of immediate behavioral relevance, analogous to how sensory attention selectively prioritizes salient features in perceptual space (31–34). This framework aligns with principles of active sensing, in which organisms continuously adjust the direction, density, and timing of information gathering to meet current task demands (35–41). Accordingly, a central prediction is that sweep modulation should emerge rapidly and flexibly across behaviors and contexts, rather than being limited to alignment with head direction or stable, experience-dependent goals.

To test this hypothesis, we performed large-scale Neuropixels recordings across the entorhinal-hippocampal circuit and upstream head-direction areas in rats engaged in behaviors with shifting attentional demands, including spontaneous pursuit of a rapidly moving bait, orienting during alert immobility, and backward locomotion, as well as during REM sleep. Across all conditions, sweeps were dynamically tuned to locations of potential behavioral relevance through modulation of three core parameters of the sweep pattern – direction, width and frequency. These findings reveal a general, attention-like mechanism that enables the navigation circuit to transition fluidly between broad, coverage-maximizing sampling and focused, behaviorally guided sampling of task-relevant locations. This flexibility indicates that hippocampal–entorhinal sweeps support a dynamic internal allocation of spatial attention that is independent of learned goals and substantially more versatile than previously recognized.

## Sweeps and internal direction are directed towards moving targets

To test whether sweeps are modulated by objects that attract the animal’s attention, we recorded neural activity from 7 rats implanted with Neuropixels probes in MEC-parasubiculum (fig. S1). Recordings were obtained while the animals foraged for randomly scattered food crumbs inside a square open-field arena (1.5 × 1.5m; Fig. 1A-B) and while they engaged, in the same environment, in a pursuit task designed to mimic natural prey-capture behavior (Fig. 1C-D). The two tasks were performed consecutively, with recordings acquired from the same neuronal populations. In the pursuit task, the rats chased a piece of food that was suspended from a string and moved rapidly and unpredictably through the arena by the experimenter (Movie S1). The fast and erratic motion of the bait required the animals to monitor it continuously and make quick course corrections on the fly. Theta sweep and internal direction patterns were compared across the two task conditions, by decoding position and direction signals from the spiking activity of many hundreds of co-recorded cells (mean of 923 cells per session), including grid cells (mean of 195 cells per session; fig. S2B) and internal direction cells (mean of 323 cells per session; fig. S2C). To extract sweeps, which in MEC are primarily expressed by grid cells (11), we decoded position from all cells at 10-ms time steps using population vector correlations, and identified coherent, outgoing trajectories within each theta cycle (11) (Fig. 1B,D and fig. S3A). To extract internal direction — which is expressed by theta-rhythmic internal direction cells in parasubiculum (11) with broad head direction tuning (16, 17) — we first used an iterative procedure to estimate each cell’s tuning to internal direction, and then used these tuning curves to decode internal direction using population vector correlations (fig. S3B).

**Fig. 1.**
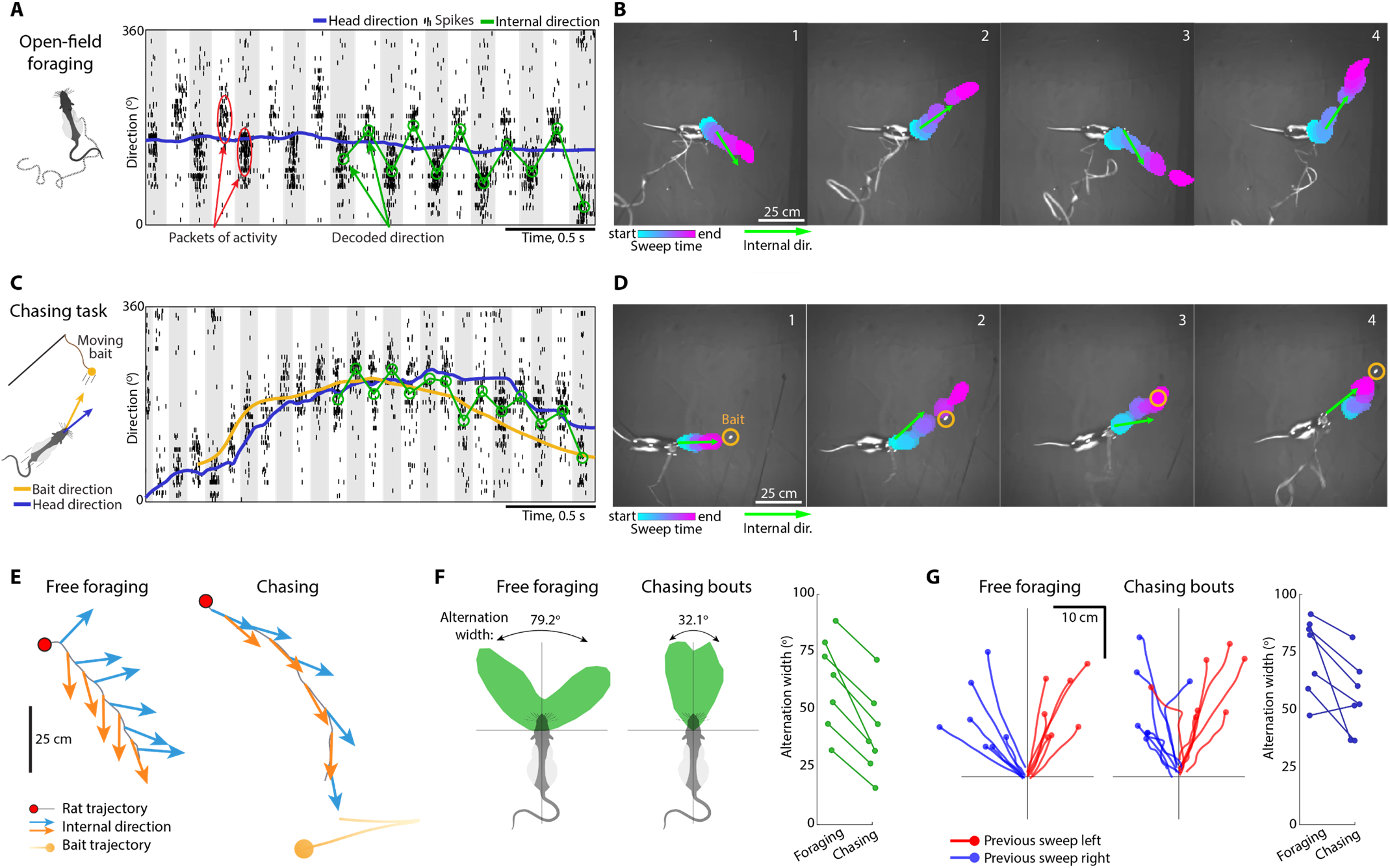
Theta-sweep are narrowly forward-focused during chasing of a moving object. (**A**-**B**) Theta sweeps during free foraging. (**A**) Left, schematic of the free foraging task. Right, raster plot with spikes from 474 internal direction cells, mostly from parasubiculum, sorted along the y-axis by each cell’s preferred direction. Note theta-rhythmic packets of population activity that alternate between left and right of the animal’s head direction (blue line) across successive theta cycles (alternating grey and white background). Green circles connected by lines show decoded internal direction for each theta cycle (decoded with reference to session-averaged population vectors as in fig. S3). (**B**) Left-right-alternating internal direction and sweeps during free foraging. Images show snapshots of the recording arena at the beginning of four successive theta cycles (top views; same session as (A)). The decoded position from the joint activity of 664 MEC-parasubiculum cells throughout each theta cycle is plotted as colored blobs, with color indicating time within the sweep. Internal direction, decoded from 474 internal direction cells, is plotted as green arrows. Note left-right-alternation of both signals with respect to the head axis. (**C**-**D**) Theta sweeps during chasing. (**C**) Left, schematic of the chasing task, where rats chase a string-suspended moving reward in the open field arena. Vectors indicate head direction (blue) and bait direction (yellow). Right, raster plot with spikes from 474 internal direction cells (same as (A)) during a bout of chasing towards the bait (plotted as (A)). Bait direction, the direction from the rat’s head to the bait, is plotted in yellow; decoded internal direction is shown in green (as in (A)). Note that while left-right alternation of population activity is maintained across theta cycles, the alternation is narrower than in the foraging task in (A). (**D**) Internal direction (green arrows) and sweeps (colored blobs) during chasing (same session as (C)). Decoded positions and directions are plotted as in (B), with bait position marked by yellow circles. Note that left-right-alternation is narrowly focused toward the bait. (**E**) Internal direction during free foraging and chasing (same recording as (A and C)). Left, decoded internal direction across 11 successive theta cycles during free foraging, plotted as blue and orange arrows (even and odd theta cycles). Note regular left-right-alternation with consistent 30-40° offsets from the head axis. Right, internal direction during a targeted run towards the bait in the chasing task. Yellow line shows the bait’s trajectory during the same time period, with a circle marking its final position. Left-right alternation is preserved, but narrowly focused forward, as the rat runs directly toward the bait. (**F**) Left panels, polar histograms of decoded internal direction (relative to head axis) during open field foraging and during bouts of chasing (same recording as in (A and C)). Note that internal direction is bimodally distributed with respect to the head axis in both conditions, but with peaks closer to the midline during chasing bouts. Alternation widths, computed as the mean angular offset in decoded direction across successive theta cycles, are indicated above. Right panel, alternation widths for internal direction during free foraging and chasing for all seven animals, one pair of dots per animal. (**G**) Left panels, averaged spatial extents of sweeps during free foraging and chasing bouts in all 7 rats, with one pair of red and blue lines per animal. Sweeps were averaged by rotating them to head-centered coordinates (with head orientation vertical) and averaging sweep position across 10-ms bins of theta cycles in which the preceding sweep went left (red) or right (blue). Note left-right alternation of sweep direction in both conditions, with sweeps more narrowly focused forwards during chasing. Right panel: Alternation widths for sweep direction during free foraging and chasing for all 7 rats, one pair of dots per animal. Alternation width was calculated as for internal direction in (F). Credit: rat (A, C and F), scidraw.io/Gil Costa.

The rats’ behavior during the chasing task was characterized by bouts of rapid chasing directly towards the bait (‘chasing bouts’), interspersed with pauses where the rat either reoriented towards the bait or disengaged from the task (fig. S2D, Movie S1). During chasing bouts, defined as periods of running straight towards the bait, internal direction and sweeps were forward-focused (Fig. 1C-E). While left-right-alternation persisted (fig. S2E), the ‘alternation width’, defined as the angular separation of internal direction in neighboring theta cycles, was narrower than during free exploration in the open field (mean ± s.e.m. of 39.8 ± 6.9° vs. 62.2 ± 7.6°; Wilcoxon signed-rank test: W = 28, n = 7, p = 0.016; Fig. 1E-F) or during running on a linear track (fig. S2G). In addition, the frequency of the theta oscillations was increased (chasing: 9.1 ± 0.2 Hz, open field foraging: 8.2 ± 0.2 Hz, Wilcoxon signed-rank test: W = 28, n = 7, p = 0.016), also at comparable running speeds (fig. S2F-H). The alternation width of internal direction signals was negatively correlated with theta frequency across behaviors (chasing, open field and linear track combined; Pearson correlation: r = –0.17 ± 0.03, mean ± s.e.m., p < 0.05 in all 7 rats; fig. S2I). A similar reduction in alternation width during chasing was observed in sweeps of grid cells (mean ± s.e.m. of 55.0 ± 6.1° during chasing vs. 74.0 ± 6.3° during free foraging, Wilcoxon signed-rank test: W = 27, n = 7, p = 0.031; Fig. 1G) and hippocampal place cells (fig. S4A). Collectively, these adjustments of the sweep pattern during chasing bouts result in focused, high-rate sampling of a narrow spatial sector between the animal and the bait.

Having seen that the width and rate of sweep sampling is dynamically modulated, we asked whether similar control is exerted over the direction, or ‘central axis’, of the sampling sector. Hypothesizing that such directional steering may be evident when the bait is not directly in front of the animal, we turned our focus to the brief pauses between chasing bouts, when rats lost track of the bait for a few hundred milliseconds and reoriented before initiating another pursuit (Fig. 2A, Movie S1). Strikingly, during these ‘reorienting events’, internal direction and sweeps often turned towards the bait before the onset of head turning, resulting in a decoupling of the neural signals away from the head axis (Fig. 2A-D, Movie S1). We quantified the presence of such ‘decoupling events’ by identifying all periods where the central axis of the internal-direction sector (green arrow in Fig. 2A) deviated more than 30 degrees from the head axis (439.6 ± 138.7 decoupling events per session, mean ± s.e.m.; 46.1 ± 3.7 events/min; Fig. 2A and E). Decoupling events were brief (410.4 ± 10.1 ms, mean ± s.e.m.), occurred during ongoing theta oscillations (fig. S5A), and coincided with periods of low movement speed. These events were often terminated by an orienting movement towards the bait, with a mean ± s.e.m. latency between peak decoupling and head turning of 84.4 ± 2.8 ms (Fig. 2E-F). During decoupling events, internal direction signals generally pointed to the same egocentric side as the bait (aligned in 73.2 ± 2.5% of theta cycles with decoupling, p<0.05 in 7/7 rats, one-tailed binomial test; fig. S5B), while left-right alternation was rare (fig. S5C). Similarly, in grid cells, sweeps were directed to the same side as the bait in 63.3 ± 1.5% of decoupling events (p<0.05 in 6/7 rats, one-tailed binomial test, fig. S5B). A similar bias was seen in hippocampal place cells (fig. S4B-C). In the majority of decoupling events recorded in internal direction cells (69.6 ± 2.7%, mean ± s.e.m.), an internal-direction-aligned orienting movement could be observed within 200 ms (Fig. 2G). In exceptional cases, the rat remained stationary for a long interval while internal direction and sweeps tracked the swinging bait (fig. S5D; Movie S2), suggesting that internal direction and sweeps do not simply reflect an imminent motor command but instead reflect allocation of spatial attention towards the target before the animal finally commits to an action.

**Fig. 2:**
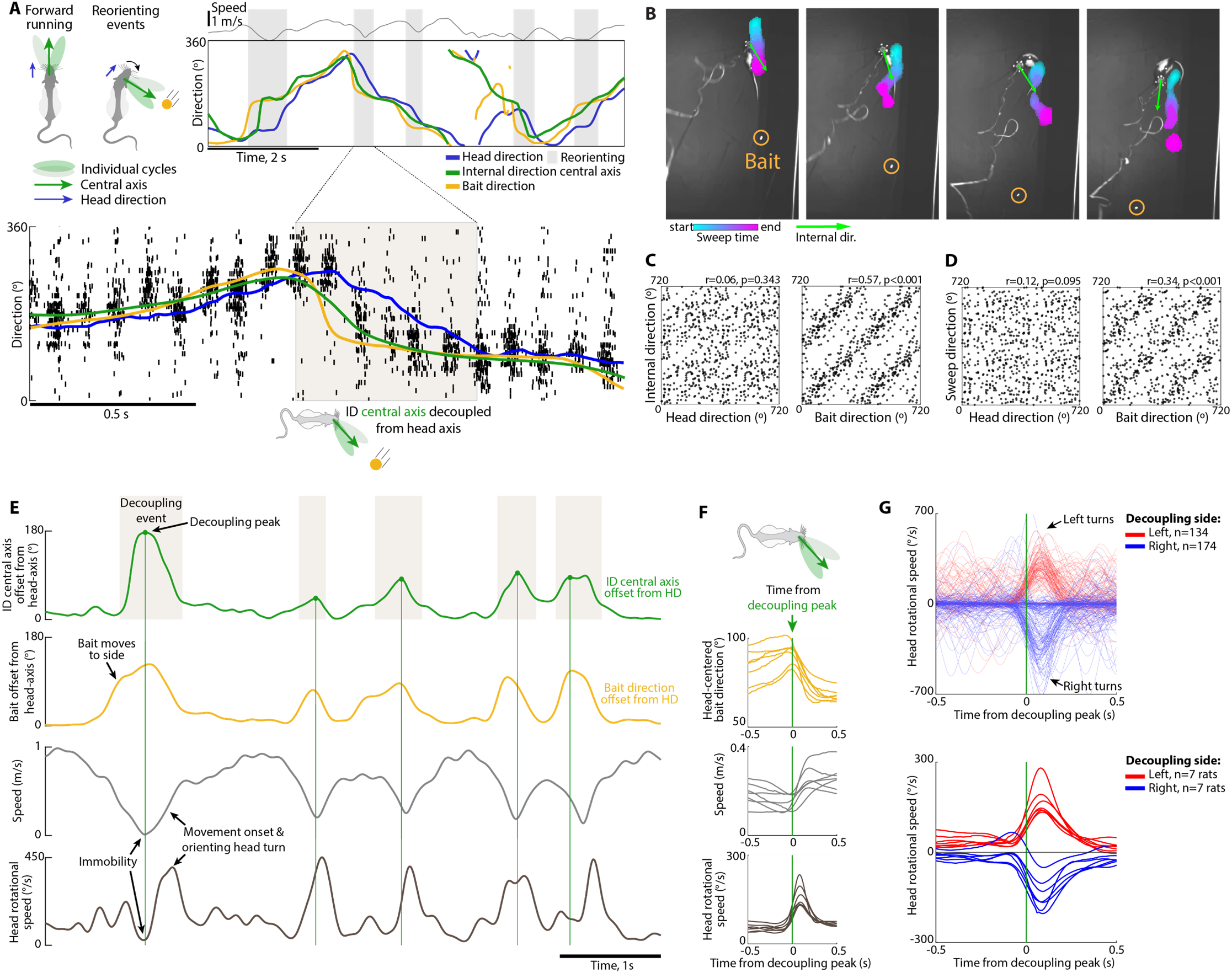
Internal direction and sweeps reorient toward a moving target before behavioral realignment. (**A**) Schematic, left-right alternation of internal direction across theta cycles (colored blobs) is centered around a slower-moving ‘central axis’ (green arrow; moving average across theta cycles) that is aligned with head direction (blue) during forward running, but may decouple from the head axis during attentional shifts, e.g. towards a bait (yellow). Top panel, tracked head direction (blue), bait direction (yellow) and running speed (top) during chasing of a moving bait (same session as Fig. 1C). The internal direction central axis, estimated by smoothing the decoded signal across theta cycles, is plotted in green. Note that periods of rapid chasing are interspersed by orienting events, where the rat stops and turns towards the bait (grey rectangles). Bottom, raster plot with spikes from internal direction cells (plotted as in Fig. 1C) during one of the orienting events (start and end indicated by straight lines). Tracked head direction, bait direction and central axis are shown. Note that during the orienting event, packets of internal-direction-cell activity are centered around bait direction, and not head direction, showing that the central axis of the internal direction signal decouples from the head axis. (**B**) Decoded internal direction and sweeps during four successive theta cycles in a reorienting event with a counterclockwise turn towards the bait (bait initially behind the rat; plotted as in Fig. 1B). Note that bait-directed internal direction signals and sweeps precede turning behavior. (**C**) Scatter plots showing alignment between the central axis of internal direction and head direction (left panel) and between internal direction and bait direction (right panel) during reorienting events in one example animal. Note that sweeps are centered around bait direction (not head direction). (**D**) As in (C), but for sweep direction. (**E**) From top: Green trace, angular offset between the central axis of internal direction (green arrow in A) and head direction; yellow trace, offset between bait direction and head direction; grey, running speed; and black, angular head speed during a 6-s segment of the chasing task (same session as in A). Note that internal direction decouples from the head axis (grey rectangles; peaks indicated by green vertical lines) whenever the bait moves to the side of the animal (yellow trace). Decoupling events coincide with immobility periods and are followed by orienting movements and initiation of new chasing bouts. (**F**) Mean head-centered bait direction (top), running speed (middle) and angular head speed (bottom) triggered by peak decoupling (green vertical line) in all seven animals, one line per animal. Decoupling of internal direction from head direction was followed by increased running speed (18.9 ± 1.3 m/s vs. 26.2 ± 1.9 m/s before and after the peak of decoupling, mean ± s.e.m.; increase of 43.3 ± 15.1%, Wilcoxon signed-rank test: W = 28, n = 7, p = 0.016), and increased angular head speed (70.5 ± 3.4 °/s vs. 142.1 ± 8.0 °/s, increase of 104.7 ± 17.0%, Wilcoxon signed-rank test: W = 28, n = 7, p = 0.016). The latency to peak acceleration was 95.7 ± 13.5 ms for running speed and 84.4 ± 2.8 ms for angular head speed, mean ± s.e.m. (**G**) Top, angular head speed triggered by decoupling peaks where internal direction pointed to the left (red; 134 events) or right (blue; 174 events) of the head axis for one example animal, each line corresponds to one decoupling peak. Note that most, but not all, decoupling peaks are followed by an ipsiversive orienting response. Bottom, mean angular head speed triggered by decoupling events where internal direction pointed to the right (blue) or left (red) for all seven animals, one pair of lines per animal. Credit: rat in (A and F), scidraw.io/Gil Costa.

## Sweeps and internal direction frequently decouple from the head axis during immobility

Having shown how internal sampling can be biased towards a moving target on a moment-to-moment basis in the pursuit task, we next asked whether sweeps and internal direction signals are similarly modulated during unconstrained behavior in the absence of an explicit goal or task. To address this question, we analyzed open-field foraging sessions (n = 10 rats, 1 session per rat), focusing on periods of immobility with preserved theta oscillations (548.9 ± 83.5 s per session, 19.0 ± 3.1% of total session duration, mean ± s.e.m. across 10 rats; fig. S6A-B). We reasoned that these alert pauses, when the animal is stationary but engaged, might expose transient shifts in the locus of covert spatial attention, analogous to the brief pauses between pursuit episodes in the bait-chasing task.

Theta-nested sweeps persisted during these immobility periods, but the rigid head-centered alternation characteristic of running was markedly reduced (Fig. 3A-B; Movie S3). The fraction of theta-cycle triplets exhibiting left-right-alternation dropped significantly compared to running (Fig. 3C-D and fig. S6C), and the central axes of internal direction and sweeps decoupled from the head axis more frequently (Fig. 3C-D). This pattern closely resembled the decoupling observed during reorienting pauses in the pursuit task (Fig. 2). Decoupling events often persisted for extended periods, lasting up to 10-20 consecutive theta cycles (Fig. 3E). During these events, sweep direction in grid cells remained aligned with internal direction rather than with head direction (Fig. 3F), a relationship that was also observed for place-cell sweeps (fig. S4D-F).

**Fig. 3.**
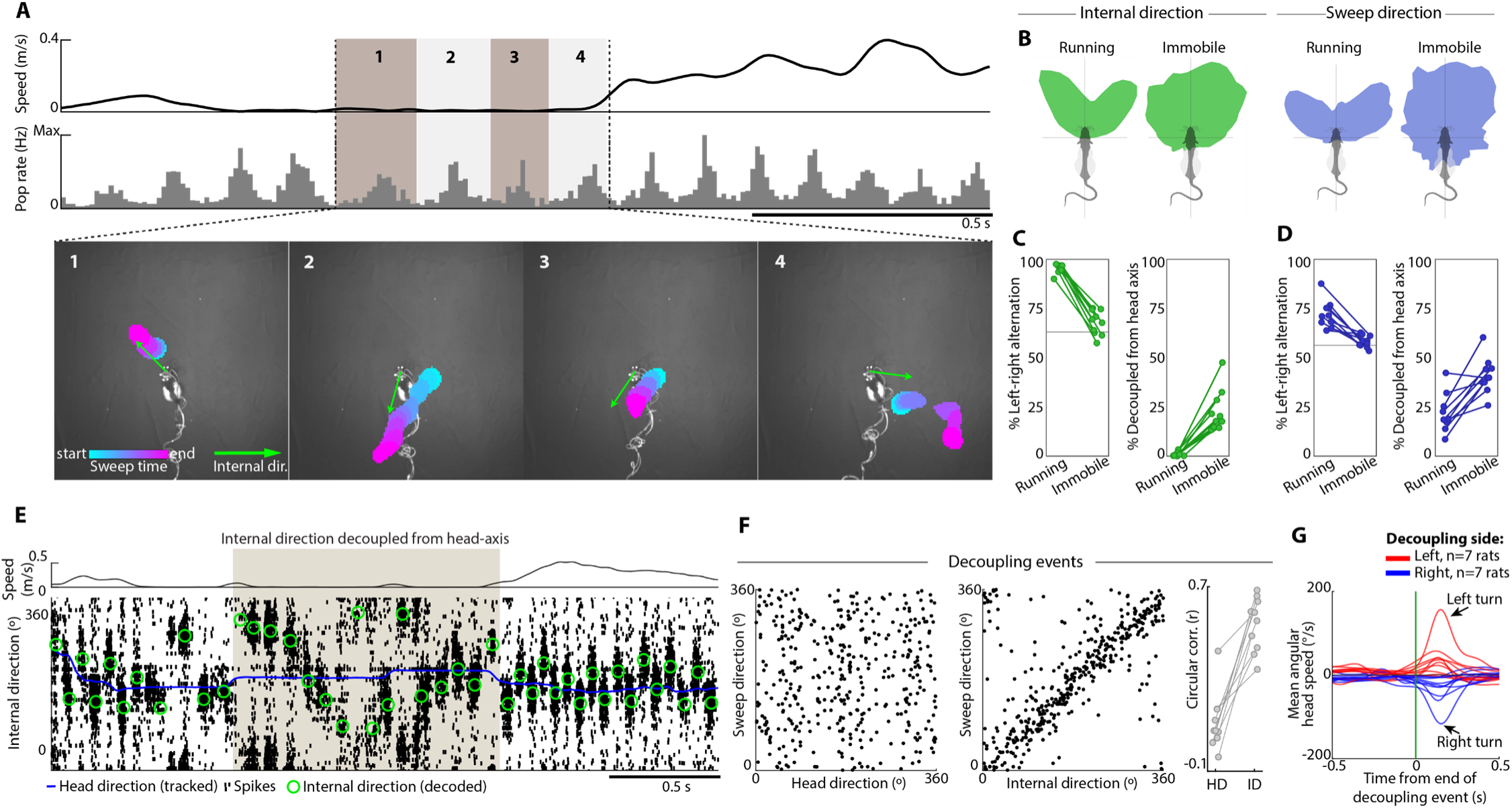
Increased variability of internal direction and sweeps during immobility. (**A**) Top, running speed during a 2-s epoch from a free foraging session. Middle, summed spike counts from 1,187 co-recorded MEC-parasubiculum cells, showing preserved 6-10 Hz theta-rhythmic population activity during immobility (first half of the epoch). Bottom: Decoded internal direction (green arrow) and position (colored blobs) during four successive theta cycles from the example in the top and middle panels (theta cycles 1-4). Internal direction and sweep signals remain aligned during immobility, but the head-centered alternation pattern is lost. (**B**) Head-centered distribution of internal direction (left panels) and sweep direction (right panels) during running and immobility periods with theta in an example open field foraging session. Note that internal direction and sweep direction is less rigidly aligned to the head axis during immobility than during running. (**C**) Left, fraction of theta-cycle triplets with left-right-alternation in the internal direction signal during running (95.4 ± 0.7%, mean ± s.e.m. across 10 rats) and immobility (67.8 ± 2.0%; Wilcoxon signed-rank test, running vs. immobility: W = 55, n = 10, p = 0.002). The fraction of cycles with alternation during immobility is near shuffled values (horizontal line; 63.2 ± 0.9%, mean ± s.e.m.). Right, fraction of theta cycles where the internal direction central axis decoupled by >30° from the head axis during running (1.1 ± 0.3°) and immobility (22.8 ± 3.4°; Wilcoxon signed-rank test: W = 0, n = 10, p=0.002). (**D**) As (C), but for sweep direction. Left-right alternation in 72.3 ± 2.2% vs. 58.3 ± 1.0% of theta-cycle triplets (shuffled: 56.2 ± 0.7%), Wilcoxon signed-rank test: W = 55, n = 10, p = 0.002. Decoupled from the head axis in 23.5 ± 2.9% vs. 41.1 ± 2.9% of theta cycles, Wilcoxon signed-rank test: W = 1, n = 10, p = 0.004. (**E**) Example of decoupling between head axis and central axis of internal direction signal during immobility. Raster shows spikes from 474 internal direction cells (sorted by preferred direction) during a period of immobility, flanked by running periods. Shaded box indicates an extended period (∼1.5 s) where decoded internal direction (green circles) deviates substantially from tracked head direction (blue line). Note that both left-right alternation and centering around head-axis are lost during this ‘decoupling event’. (**F**) Left panels, scatter plots of sweep direction vs. tracked head direction (left) or internal direction (right), confined to immobility periods where internal direction is decoupled from head direction (data from one animal, one session). Note that sweeps are aligned to internal direction, and not head direction, during these events. Right, circular correlation coefficients between sweep direction and head direction (r = 0.11 ± 0.04, p < 0.01 in 3/10 rats) or internal direction (r = 0.54 ± 0.04, p < 0.01 in 10/10 rats) for all 10 rats, one pair of points per rat. Wilcoxon signed-rank test: W = 55, p = 0.002, n = 10. (**G**) Mean angular head speed triggered by the end-times of decoupling events during immobility periods in the open-field task where internal direction pointed to the right (blue) or left (red) for all ten animals, one pair of lines per animal. Note that decoupling events are followed by an ipsiversive orienting response in the averaged traces of all animals (mean ipsiversive rotational speed of 4.4 ± 1.7 °/s vs. 26.7 ± 8.3 °/s before and after end of decoupling events, respectively, mean ± s.e.m. across 10 rats; Wilcoxon signed-rank test: W = 49, n = 10, p = 0.027). Credit: rat in (B), scidraw.io/Gil Costa.

Decoupling events during free foraging sessions were often followed by movement, parallelling observations in the chasing task (fig. S6D). Overall, 33.8 ± 1.5% (mean ± s.e.m.) of internal-direction decoupling events were followed by a head turn within 200 ms, and the majority of these turns (65.1 ± 3.8%) were aligned with the internal direction signal decoded at the end of the decoupling event (Fig. 3G). Post-hoc inspection of video recordings revealed that many decoupling events were targeted towards transient events such as incoming food crumbs (Movie S3).

Taken together, these findings show that the sweep system remains active during immobility (25), but with looser coupling to the head axis and with less rigid alternation dynamics, enabling focused but flexible sampling of surrounding space that often precedes orientating movements and subsequent locomotion.

## Sweeps and internal direction track movement direction when movement and head direction are in conflict

The brief shifts in sweeps and internal direction signals towards moving targets observed here mirror the reflexive allocation of spatial attention to salient stimuli (42, 43). However, spatial attention is guided not only by salient external events but also by internally generated movement plans (42–44). To test whether sweeps align with the direction of impending movement in the absence of salient stimuli – and independently of head direction or reward location – we examined conditions in which these variables could be dissociated for extended periods. As one such condition, we examined sweep dynamics during backward locomotion, which we induced using two complementary paradigms: guiding rats into a narrow, dead-ended corridor with reward at the end, requiring them to back out (n = 2 rats; Fig. 4A; Movie S4), and having rats drag a piece of food backward while the experimenter applied opposing force (n = 3 rats; fig. S7C; Movie S5). These designs allowed us to determine whether sweep direction follows the direction of impending movement rather than the orientation of the head or the location of reward sites.

**Fig. 4.**
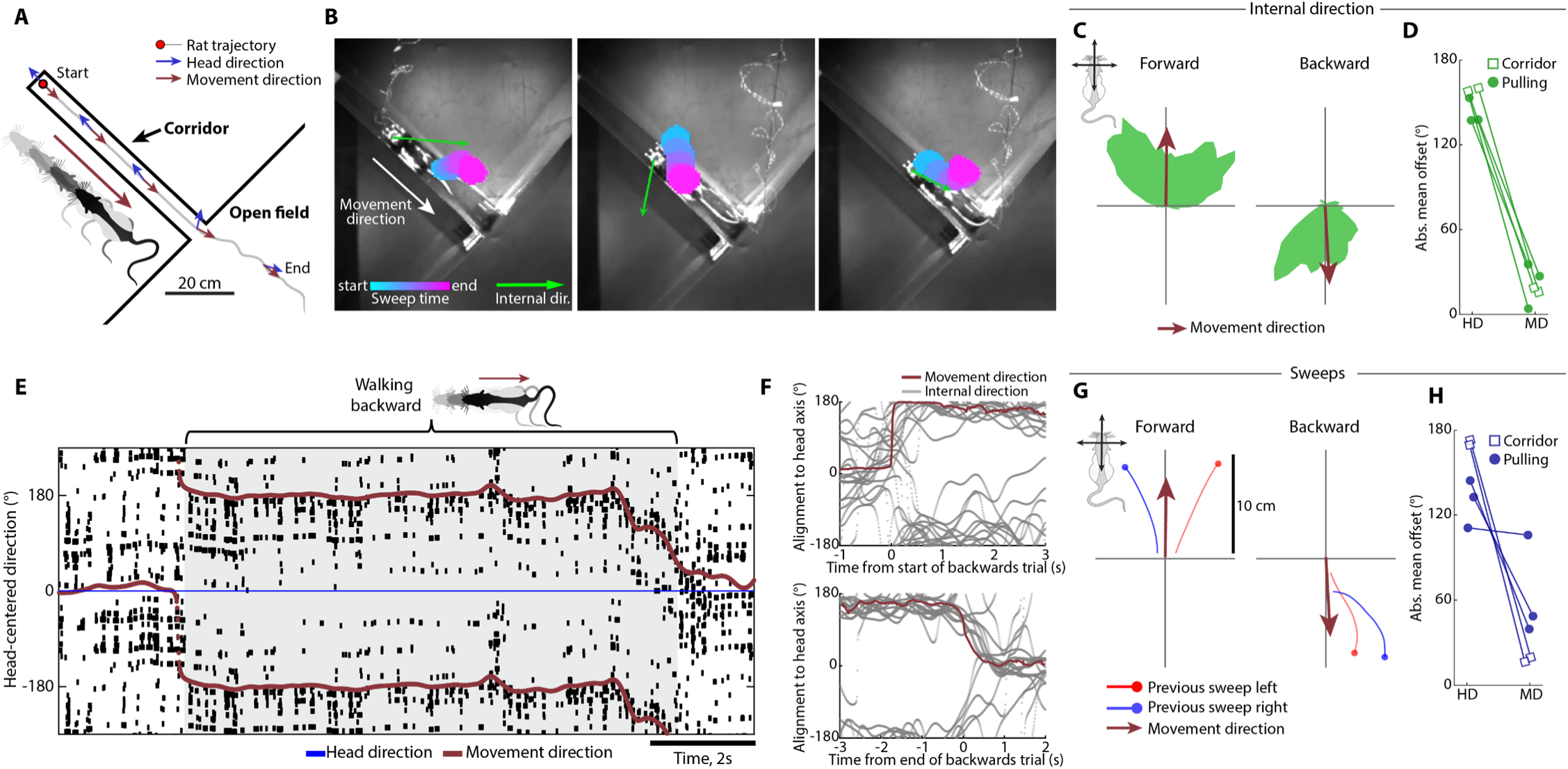
Sweep direction aligns with movement direction during backward locomotion. (**A**) Schematic of the backward-walking task, where rats backed out of a narrow, dead-ended corridor. Grey line shows the rat’s trajectory out of the corridor during one of the trials. Arrows show tracked head– and movement direction at regular intervals throughout the trial. (**B**) Decoded sweeps and internal direction during three example theta cycles as a rat backs out of the corridor (plotted as in Fig. 1B). Note that both sweeps and internal direction point backwards. (**C**) Polar histograms showing head-centered distribution of decoded internal direction during forwards (left) and backward locomotion (right) in an example recording session (same session as (B)). Brown arrow shows mean movement direction. (**D**) Alignment of internal direction to head direction (HD) vs. movement direction (MD), quantified as the absolute mean angular offset from HD or MD during periods of backward movement across animals (n = 5 rats, one pair of points per rat). Symbols indicate whether rats were tested in the corridor task (open rectangles) or the backward-pulling task (filled circles). Internal direction is aligned to MD but not HD (Wilcoxon signed-rank test, one-sided (MD offset < HD offset): W = 0, p = 0.031, n = 5). (**E**) Raster plot of spikes from 121 internal direction cells during a 9 s segment of backward walking (shaded rectangle). Spikes are sorted along the y-axis based on the cells’ preferred allocentric direction and centered around the animal’s head axis (forwards: 0°, backwards: 180°). Movement direction (relative to head axis) is plotted in brown. Y-axis is repeated beyond 180° for visualization. Note that the internal direction signal points backwards in alignment with movement direction throughout the trial. (**F**) Central axis of internal direction (head-centered, smoothed across theta cycles) during all backwards trials in one animal, aligned to the onset of backward movement (top) and the offset of backward movement (bottom). Mean movement direction (head-centered) is shown in brown. Note rapid and persistent inversion of the central axis as movement direction changes. (**G**) Sweeps, from grid cells in an example session, rotated to head-centered coordinates (head orientation is vertical) and averaged across theta cycles during periods of forward (left panel) or backward locomotion (right panel), for theta cycles where the preceding sweep went left (red) or right (blue). Note that sweeps are directed backwards during backward movement. **(H)** Alignment of sweep direction to head direction (HD) vs. movement direction (MD), as in (D), during periods of backward movement (n = 5 rats, 1 session per rat). Sweep direction is generally aligned to MD and not HD (Wilcoxon signed-rank test, one-sided (MD offset < HD offset): W = 0, p = 0.031, n = 5). Credit: rat in (A, C and G), scidraw.io/Gil Costa.

We first asked whether sweep dynamics would realign to the direction of movement even when that direction was opposite to the head axis and uninformative about reward. The corridor task produced extended bouts of uninterrupted backward locomotion along a fixed axis, whereas the pulling task generated backward trajectories spanning multiple allocentric directions. Although theta oscillations during backward walking were slower and lower in amplitude than during forward running (fig S7A-B), both sweeps and internal direction signals remained robustly expressed across paradigms (Fig. 4B; fig S7C). Strikingly, at the onset of backward locomotion, the central axis of internal direction rotated by 166.3 ± 11.7° (mean ± s.e.m.) relative to the head axis and then remained aligned with the movement vector, with a residual offset of only 5.3 ± 8.7° across tasks (Fig. 4C-F). This inversion occurred rapidly around the onset of backwards movement and persisted throughout the backward-walking period (Fig. 4E-F; fig. S7D-E). A comparable rotation was observed in sweeps of grid cells: their central axis shifted by 151.5 ± 12.1° relative to head direction and by 36.4 ± 16.8° relative to movement direction (mean ± s.e.m. across both tasks; Fig. 4G-H). Left-right-alternation was reduced during backward walking, occurring in 63.2 ± 3.3% of theta-cycle triplets for internal direction (shuffled: 54.3 ± 2.2%) and in 56.4 ± 1.7% of triplets for sweeps (shuffled: 51.3 ± 0.4; fig. S7F-G). Backward-pointing sweeps were also evident in hippocampal place cells (fig. S4G-I). Finally, during passive displacement by the experimenter (e.g., when rats were lifted into and out of the arena), internal direction signals frequently rotated to align with the imposed movement vector rather than the head axis (Movie S5), indicating that the sweep system tracks movement vectors independently of how displacement is generated.

Collectively, these observations demonstrate that sweeps and internal direction are governed not only by salient external stimuli or explicit rewards but also by internally generated movement plans. In this sense, the sweep system provides a unified mechanism through which both exogeneous and endogenous control signals can bias the allocation of spatial attention on a moment-to-moment basis – mirroring the well-established division between reflexive and volitional attention in humans and non-human primates (34, 42, 45).

## Variability in sweep patterns is preserved during REM sleep

The backward locomotion task showed that sweeps can be guided by internal mechanisms such as planned movement direction. To test whether overt movement is required for such internally driven modulation, we next turned to rapid eye movement (REM) sleep, a state that provides a natural assay of intrinsic network dynamics under conditions when external sensory input and motor output are largely suppressed (11, 46–48). Using decoding analyses, we found that the sweep system remains robustly active during REM sleep and continues to switch between “exploratory” and “focused” sampling modes, indicating that these dynamics can be generated endogenously, independent of overt behavior.

We obtained sleep recordings from 7 rats with MEC-parasubiculum implants and identified periods of REM sleep (18.7 ± 3.2 minutes of REM sleep per session, mean ± s.e.m.) based on electrophysiological and behavioral criteria. We first investigated the coherence of internal direction and position signals across entire REM sleep episodes, using tuning curves from a same-day open-field foraging session as a reference for decoding (6-22 REM sleep episodes per rat; mean episode duration: 78.2 ± 9.0 s; Fig. 5A; fig. S8A-D). Throughout REM sleep, population activity remained internally consistent, forming a bump of activity that traversed the population with temporal dynamics resembling physical movement in the awake state (fig. S8A-B). In grid cells, decoded trajectories often spanned several meters in the open-field reference space over the course of a REM episode (Fig. 5A). Co-recorded grid modules expressed highly similar spatial trajectories (fig. S8C; trajectory correlation: r = 0.47 ± 0.03, mean ± s.e.m.; p < 0.001 in 8/8 module pairs from 4 animals with >40 co-recorded grid cells in at least two grid modules). Decoding at sub-theta timescales revealed that left-right-alternating internal direction signals and sweeps were nested atop these slower, behavioral-timescale trajectories (Fig. 5A, fig. S8E-G)(11).

**Fig. 5.**
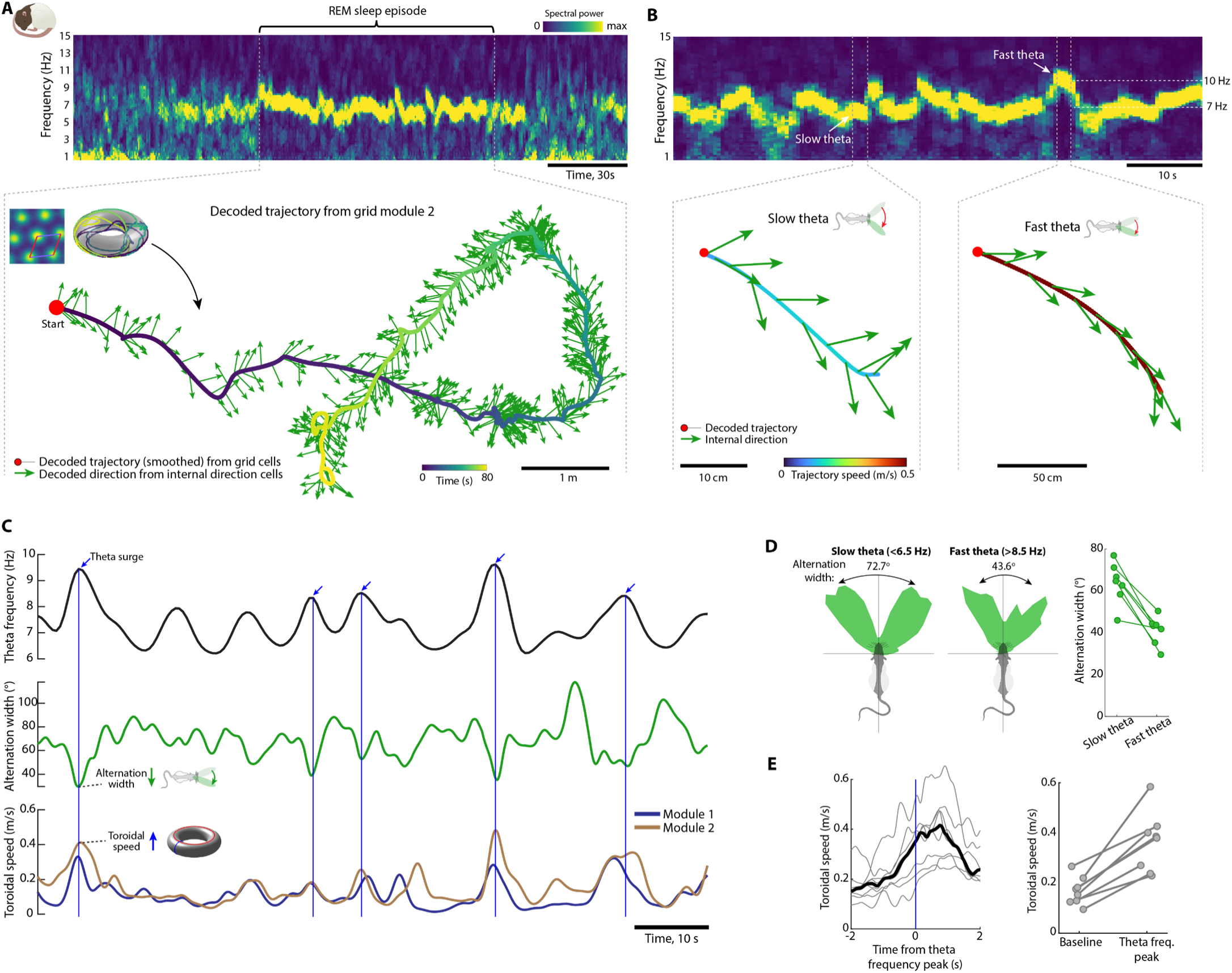
Forward-focused directional sampling-during phasic REM sleep. (**A**) Extended spatial trajectories and left-right alternation during REM sleep. Top, spectrogram of MEC-parasubiculum multi-unit activity during a ∼3-minute period of sleep. Note prominent 6-7 Hz theta oscillations during an 80-s segment of REM sleep (start and end indicated by dashed lines). Bottom, decoded trajectory during the REM sleep segment in the top panel. Based on activity of grid cells from grid module 2 in the open field (123 cells, mean spacing of 103 cm), a trajectory is decoded onto the toroidal grid-cell manifold (inset) and unwrapped into 2D space. The trajectory is color-coded by time. Decoded internal direction (from internal direction cells) is plotted as arrows (one per theta cycle). (**B**) Theta-frequency-dependent modulation of left-right alternation in internal direction cells and movement on the grid-cell torus during REM sleep. Top, spectrogram of MEC-parasubiculum multi-unit activity during a 1-min period of REM sleep. Note prominent 6-7 Hz theta oscillations with occasional “theta surge events” where frequency accelerates to ∼10 Hz (visible also in (A)). Bottom, decoded spatial trajectory from grid module 2 (solid line, color-coded by toroidal speed) and decoded direction from internal direction cells (green arrows) during snippets of slow theta (left) and fast theta (right) extracted from the top segment (between pairs of dashed lines). Note that, during the fast theta events, left-right alternation is narrowly forward-focused and toroidal speed is increased, mirroring the findings from the wake state (see Fig. 1, fig. S2). (**C**) From top: theta frequency, internal direction alternation width and toroidal speed during the 1-min segment of REM sleep shown in (B). Note that peaks in theta frequency (blue arrows) coincide with a narrowing of alternation width, decoded from internal direction cells (middle), and with increased toroidal speed, decoded from grid cells (bottom). (**D**) Left panels, histograms of decoded internal direction during REM sleep epochs with slow (left) or fast theta oscillations (right). Decoded direction is referenced to the moving-average, or central axis, of decoded internal direction (vertical axis), which serves as a proxy for head direction during REM sleep. Note that alternation width is narrower during REM sleep with fast theta, compared to REM sleep with slow theta. Right panel, alternation widths for internal direction during REM sleep with slow and fast theta for all seven animals. (**E**) Left, speed of decoded trajectory on the toroidal manifold of individual grid modules, triggered by peaks in theta frequency (blue vertical line). Data from 8 modules in 5 animals with >10 theta surge events are shown (grey lines). Solid black line shows mean across modules. Note peak in toroidal speed immediately after the theta rhythm accelerates. Right, mean toroidal speed during one-second periods before (baseline: t: –2 to –1 s) and immediately after (t: 0 to 1 s) theta frequency peaks. Credit: rat in (A-D), scidraw.io/Antonis Asiminas (A) and Gil Costa (B-D).

Given these observations, we asked whether the relationships between sweep dynamics, theta frequency, and running speed observed during wake persist in REM sleep. During wake, we observed a switch from wide-angled ‘exploratory’ sampling during foraging, to a narrow ‘focused’ mode during chasing and orienting movements, with accompanying surges in theta frequency during chasing bouts (Fig. 1, fig S2). Sweep dynamics varied systematically with theta frequency during REM sleep as well (Fig. 5B). Robust theta oscillations were expressed at 6-7 Hz (fig. S8H), interspersed with bouts of intense spiking activity in which theta frequency accelerated to ∼10 Hz (‘theta surges’; Fig. 5B-C). These slow and fast theta periods resemble ‘phasic’ and ‘tonic’ REM sleep substates (49, 50). During REM sleep epochs with slow theta oscillations (<6.5 Hz), internal direction exhibited wide left-right alternation (alternation width: 63.7 ± 3.7°, mean ± s.e.m, n = 7 rats), whereas fast theta epochs (>8.5 Hz) were associated with narrow alternation patterns (41.2 ± 2.6°; Fig. 5B-D). Alternation width was negatively correlated with theta frequency during REM sleep in 4 out of 7 animals (correlation: r = –0.11 ± 0.03, p < 0.01 in 4/7 rats; fig. S8I).

To test whether fast theta periods during REM sleep correspond to segments of rapid movement on the toroidal manifold, we decoded position from grid cells and measured the speed of the decoded trajectory (Fig. 5E). Grid modules with >40 cells in sessions containing >10 theta surges were included for analysis (n = 8 grid modules from 5 rats). Toroidal speed doubled during surges in theta frequency (219.7 ± 13.3% of baseline speed, mean ± s.e.m.), reaching an average of 36.4 ± 4.2 cm/s (mean ± s.e.m.) during the first second after the theta frequency peak (Fig. 5E). These findings mirror the relationship between theta frequency, alternation width and running speed during wake exploration and chasing behavior (Fig. 1, fig. S2).

Taken together, these results suggest that the brain’s navigation circuit simulates a repertoire of wake-like behavioral trajectories during REM sleep, with internal direction and sweeps sampling space according to the same adaptive logic used during wake. Sweep modulation can thus be driven by entirely endogenous mechanisms, extending the internally driven control observed during backward locomotion into a state devoid of overt behavior.

## Directional alternation and decoupling from head axis are implemented downstream of classical head direction cells

The observation that the central axis of the internal direction signal is not fixed to the head, but can shift dynamically towards targets and movement trajectories, raises the question of how these signals relate to the canonical head-direction representation in brain areas that project to parasubiculum. To determine whether shifts in the central axis of internal direction are reflected in upstream head direction signals, we recorded spiking activity from head direction (HD) cells in anterodorsal thalamus, dorsal presubiculum and postrhinal cortex, along with MEC-parasubiculum cells, using up to four chronically implanted Neuropixels probes and recording between 1,161 and 2,975 cells simultaneously across areas (n = 6 rats; Fig. 6A and fig. S1 and S9). Decoding analysis revealed that, unlike simultaneously recorded internal direction cells in parasubiculum, head direction signals in anterodorsal thalamus (38-109 HD cells per session), dorsal presubiculum (194-250 HD cells per session) and postrhinal cortex (16-136 HD cells per session) remained tightly aligned to the animal’s head axis throughout the session, without detectable left-right alternation at the theta timescale (Fig. 6B, fig. S9A-H). In these regions, alternation was identified, respectively, in 61.4 ± 0.9%, 62.6 ± 0.7% and 64.2 ± 0.7% of theta-cycle triplets during running (mean ± s.e.m. for presubiculum (3 rats), anterodorsal thalamus (2 rats) and postrhinal cortex (3 rats), all comparable to chance level (mean shuffled values of 65.2 ± 0.3%). In contrast, in parasubiculum, alternation was observed in 92.2 ± 1.0% of theta cycles in the same sessions in the same rats. Importantly, in the chasing task (n = 4 rats), when the parasubicular internal direction signal decoupled from the head axis and pointed towards the bait, thalamic and presubicular population signals remained aligned to head direction, rather than bait direction (Fig. 6B-D; absolute offset from head direction: 20.3 ± 5.3°, offset from bait direction: 83.7 ± 9.7°, mean ± s.e.m. across 5 samples of decoded direction from anterodorsal thalamus and presubiculum from 4 rats, 1 session per rat; Wilcoxon signed-rank test, left-sided: W = 0, n = 5, p = 0.031). Similarly, during backward walking (n = 3 rats), head direction cells in anterodorsal thalamus and presubiculum remained aligned with head direction, rather than movement direction (Fig. 6E-H, mean offsets of 10.3 ± 2.8° and 154.2 ± 9.3° from head direction and movement direction, mean ± s.e.m. across 5 samples of decoded direction from anterodorsal thalamus and presubiculum in 4 sessions from 3 rats; Wilcoxon signed-rank test: W = 0, n = 5, p = 0.031).

**Fig. 6.**
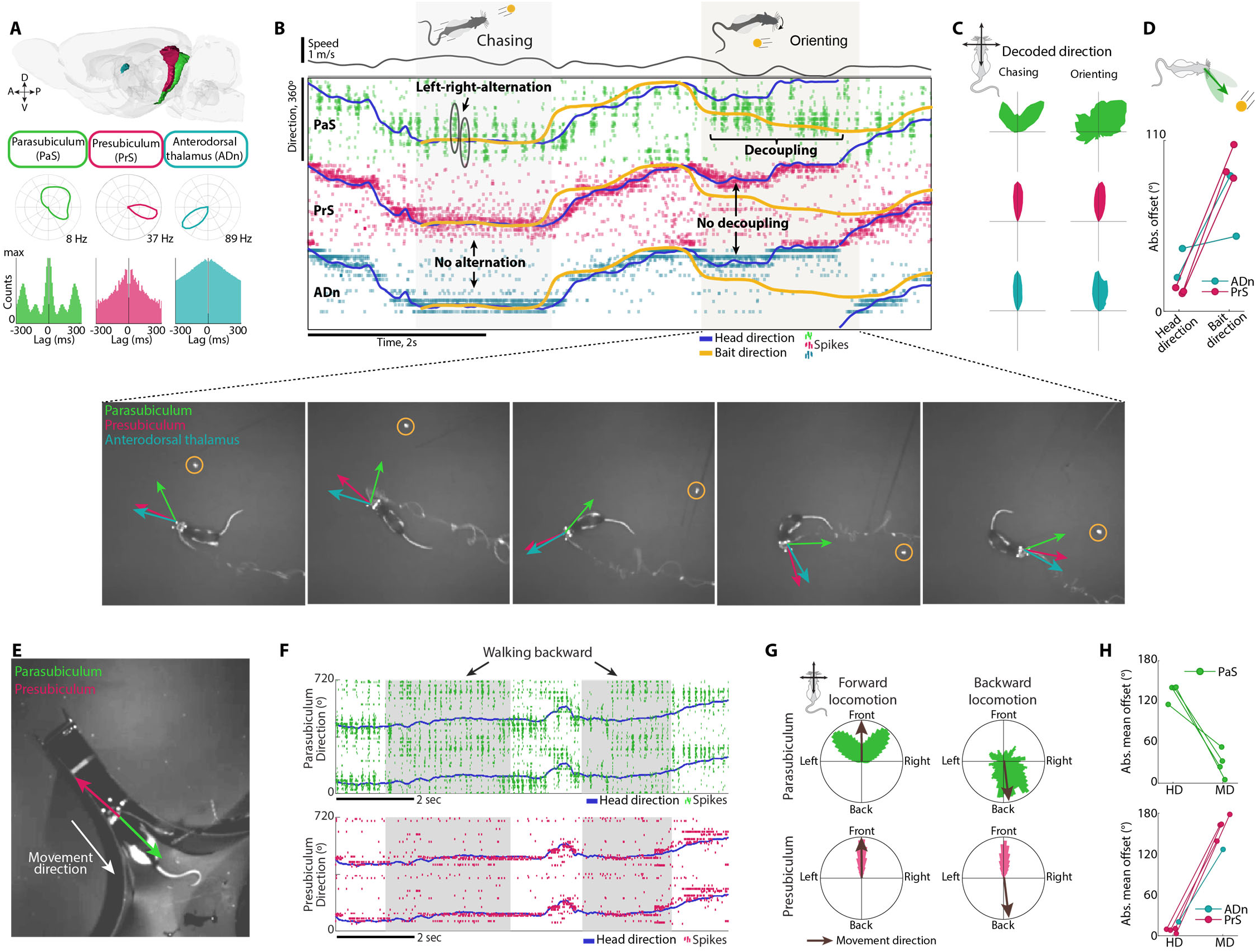
Upstream head-direction cells provide a stable allocentric reference during decoupling events. (**A**) Regional differences in directional tuning. Top row, three key nodes of the head-direction circuit: parasubiculum (PaS, green), presubiculum (PrS, magenta) and anterodorsal thalamus (ADn, teal). Head direction tuning curves (middle row) and firing-rate autocorrelograms (bottom row) for three example cells are shown. Note broad directional tuning and theta-skipping in parasubiculum, while cells in anterodorsal thalamus and presubiculum are sharply tuned and non-rhythmic. (**B**) Top, raster plots showing spikes from direction-tuned cells in all three regions recorded simultaneously with four Neuropixels 2.0 probes in the chasing task. Units are sorted by preferred direction and brain region (PaS, 188 units; PrS, 205 units; ADn, 18 units). Tracked head direction is plotted in black and bait direction in yellow. Note that population activity in anterodorsal thalamus and presubiculum is sharply centered around tracked head direction during bouts of chasing (first shaded rectangle) as well as during orienting events (second shaded rectangle), with no signs of left-right alternation or decoupling from the head axis. Bottom, decoded direction signals from all three regions during five evenly spaced snapshots extracted from the orienting event in the top panel. Note alignment with actual head direction in presubiculum and anterodorsal thalamus. (**C**) Polar histograms showing head-centered distribution of decoded direction signals (same animal and session as in (B)) from each of the three regions (rows) during chasing bouts (left column) and orienting events identified as breaks between chasing bouts (right column). Note precise head-direction representations in anterodorsal thalamus and presubiculum across conditions. (**D**) Absolute angular offset between decoded direction from anterodorsal thalamus or presubiculum and either head direction or bait direction during decoupling events where internal direction (decoded from parasubiculum) points towards the bait. Data from 5 cell samples from 4 animals are shown. Note that anterodorsal and presubicular signals remain aligned to head direction (offset of 20.3 ± 5.3°, mean ± s.e.m.) rather than bait direction (offset 83.7 ± 9.7°; Wilcoxon signed-rank test: W = 0, n = 5, p = 0.031). (**E**) Decoded direction signals from parasubiculum (green) and presubiculum (magenta) during backward walking in a narrow corridor. Symbols as in Fig. 3B. (**F**) Raster plots showing spikes from direction-tuned cells in parasubiculum (top; 206 units) and presubiculum (bottom; 63 units) during a 10-s segment with two periods of backward walking (shaded rectangles) in the bait-pulling task. Symbols as in (B). Note that the parasubicular internal direction signal points backward, while the presubicular direction signal remains aligned with head direction. (**G**) Polar histograms showing head-centered distribution of decoded direction from parasubiculum (top row) and presubiculum (bottom row) during all segments of forward locomotion (left column) and backward locomotion (right column) in one example session from one rat (same session as (F)). Mean movement direction is shown by a brown arrow. (**H**) Top, offset between decoded direction from parasubiculum and head direction (HD) or movement direction (MD) during all backwards events in 3 rats. Bottom, same as top, but with direction decoded from anterodorsal thalamus or presubiculum (n = 5 region-specific cell samples from 4 sessions in 3 rats). Note that during backward walking, upstream head direction signals are closely aligned to head direction but not movement direction (mean ± s.e.m. offsets of 10.3 ± 2.8° and 154.2 ± 9.3° for HD and MD respectively; Wilcoxon signed-rank test: W = 0, n = 5, p = 0.031). Credit: rat in (B-D and (G), scidraw.io/Gil Cos

Finally, we investigated the coordination between parasubicular internal direction signals and upstream head-direction signals during REM sleep (n = 4 rats). Internal direction signals were mostly in register with upstream head direction signals during REM sleep (fig. S9I-J)(*51*), with left-right-alternation centered around the head-direction signal (fig. S9K). However, we also identified brief periods where internal direction signals decoupled from upstream head-direction cells (fig. S9L), resembling decoupling events during wake (Fig. 2-4), suggesting again that decoupling from the head axis is implemented downstream of the canonical head-direction circuit, within parasubiculum and MEC. This hierarchical organization allows the navigation circuit to maintain a stable allocentric reference signal in anterodorsal thalamus and presubiculum, while parasubiculum introduces a dynamic component that can bias internal sampling toward salient targets, perhaps by way of coordinate-transformation principles seen in the *drosophila* central complex (*52–55*).

## Discussion

We show that the entorhinal-hippocampal sweep system exhibits multiple defining properties of a spatial attention mechanism. In its default mode, the canonical left-right alternation of sweeps samples a broad sector of space ahead of the animal. By contrast, during behaviors with directed spatial attention, the direction and width of this sector are flexibly retuned to prioritize momentarily relevant locations. During pursuit of a moving target, the sweep sector narrows while theta frequency increases, resulting in focused, high-rate sampling of the space between the animal and the target. When the animal temporarily loses and regains track of the target, sweeps reorient towards the displaced bait before any overt movement. This stimulus-driven redirection of sweeps is complemented by internally generated modulations: during backward locomotion, sweeps reverse along the movement axis, and during REM sleep, periods of fast theta narrow the sampling sector in the absence of external stimuli or overt movement. Together, these findings identify sweeps as a candidate neural substrate for covert allocentric spatial attention during navigation.

This attention-based interpretation provides a unifying framework for the present observations and prior reports of goal-directed hippocampal sweeps that emerge with learning (9, 10, 22–27). In those studies, remembered goal locations were expressed either as biases in session-averaged sweep trajectories (23, 24, 27) or as intermittent goal-directed events interspersed among left-right-alternating sweeps (26). By resolving both behavior and neural population activity at millisecond timescales, we show here that sweep dynamics are continuously regulated and flexibly deployed across a wide range of behaviors, without requiring prior spatial learning or an explicit reward structure. The spontaneous tracking of a rapidly moving object, the immediate alignment of sweeps with movement direction during backward locomotion, and the flexible scanning of surrounding space during immobility, all argue against an interpretation in which sweeps merely signal static, learned goal locations. Instead, our observations support a broader role for theta sweeps in dynamically selecting allocentric locations for enhanced internal processing based on moment-to-moment behavioral relevance.

The behavior-related modulation of sweeps parallels core features of active sensing, in which information sampling is directed towards attended regions of space (37, 39, 56). A particularly striking similarity to the sweep system is seen in echolocating bats, which dynamically adjust call rate, beam direction, and angular separation of sonar emissions during navigation and target pursuit (40, 41, 57, 58), mirroring the modulation of sweep frequency, central axis, and left-right alternation width observed in this study. This shared set of adjustable parameters places theta sweeps within a broad class of active sensing systems that alternate between exploratory regimes that maximize spatial coverage and focused regimes that concentrate sampling on behaviorally relevant targets (35–37, 57, 59). By leveraging similar control principles, theta sweeps may function as an internalized active-sensing mechanism that operates within the cognitive map rather than through peripheral sensors.

The covert and selective sampling of locations within entorhinal-hippocampal maps also parallels spatial attention in sensory systems, where neuronal gain is selectively enhanced and spatial tuning sharpened for behaviorally relevant locations in egocentric space, regardless of whether or not it is followed by movement (31–34). In such systems, covert attention is often conceptualized as a mental spotlight that rhythmically samples the sensory environment (60–62), a characterization that aligns naturally with the periodic structure of theta sweeps. By way of sweeps, grid cells and place cells may rhythmically index attended locations in the cognitive map, in addition to representing self-position. This framework offers a potential explanation for why hippocampal firing in highly visual species that survey their environment from a distance—such as primates (63–65) and birds (66)—is often more tightly coupled to viewed locations than to physical position. By acting as self-referenced vectors within the allocentric hippocampal map, sweeps may thus provide a functional bridge between attentional selection in egocentric sensory coordinates and allocentric spatial representations.

Finally, our findings implicate the parasubiculum as a candidate control node for dynamically steering entorhinal-hippocampal sweeps toward behaviorally relevant regions of space. Notably, the observation that parasubicular internal direction signals can decouple from upstream head direction suggests that additional inputs are integrated with head-direction information within parasubiculum to flexibly set the orientation of each sweep. The resulting internal-direction signal could then be conveyed to grid cells and, in turn, to place cells, with key features of the sweep pattern – its central axis, left-right alternation width, and frequency – already specified at the level of parasubiculum. This organization is consistent with a model in which parasubiculum exerts moment-to-moment top-down control over spatial sampling within the cognitive map, rather than merely relaying a static directional reference. Although the circuit mechanisms governing individual sweep parameters remain to be established, relevant principles may be drawn from coordinate-transforming circuits in the fly central complex (52–55), as well as from work on left-right alternation and frequency control in central pattern generator systems (67–71).

Taken together, our observations reveal a common organizational principle underlying sweep dynamics. Across diverse behaviors, we find that theta sweeps operate as a flexible, internally generated spatial sampling mechanism whose properties are continuously tuned to moment-to-moment behavioral relevance. These findings suggest that sweeps do not merely reflect navigation-related computations or learned goal representations, but instead constitute a general-purpose mechanism for directing allocentric attention within the cognitive map. By rhythmically selecting locations for enhanced processing, sweeps may provide a bridge between sensory-attentional selection and hippocampal spatial representation, offering a principled account of how internal models can be queried, updated, and steered in real time. This perspective reframes the sweep system as an intrinsic attentional infrastructure embedded within the entorhinal–hippocampal circuit—one capable of dynamically shaping the flow of spatial information to support adaptive behavior.

## Acknowledgments

We thank R.J. Gardner for development of preprocessing pipelines; S. Ball, K. Haugen, D.J. Hayden, E.H. Holmberg, K.J. Jenssen, E. Kråkvik, and H. Waade for technical assistance; the veterinary staff for animal care; M.P. Witter for advice on implantation coordinates and assessment of recording locations; and R.J. Gardner, B.R. Kanter, M. Guardamagna and R. Saxena for discussions.

## Funding

European Research Council Synergy Grant 951319 (EIM)

Research Council of Norway: Centre of Neural Computation 223262 (EIM, MBM)

Research Council of Norway: Centre for Algorithms in the Cortex 332640 (EIM, MBM)

Research Council of Norway National Infrastructure grant (NORBRAIN, 295721 and 350201).

Kavli Foundation

Ministry of Science and Education, Norway (EIM, MBM)

Faculty of Medicine and Health Sciences (AZV)

## Author contributions

All authors contributed to conceptualization and planning of experiments and analyses; AZV and MFS implanted animals and performed experiments; AZV wrote software; AZV curated, analyzed and visualized data with input from all authors; all authors contributed to interpretation; AZV and EIM wrote the paper, with periodic inputs from MFS and MBM. MBM and EIM supervised and funded the project.

## Data, code, and materials availability

The datasets generated during the current study and the code for reproducing the analyses will be available before publication.

## Supplementary Materials

Materials and Methods

Figs. 1 to 6

Figs. S1 to S9

Movies S1 to S5

## Materials and Methods

### Subjects

Data was collected from 10 Long Evans rats (3 females, 7 males; age of 10-20 weeks and weight of ∼250-500 g at time of surgery). Some of the MEC-parasubiculum data from 8 of these rats have been used for other analyses in published studies (*11*, *48*). The rats were housed together with their littermates before surgery and were thereafter singly housed in large, enriched metal cages (95 × 63 × 61 cm) or smaller Plexiglas cages (45 × 44 × 30 cm). They were kept on a 12-h light–12-h dark schedule in humidity and temperature-controlled rooms. Experiments were approved by the Norwegian Food Safety Authority (FOTS ID 18011 and 29893) and performed in accordance with the Norwegian Animal Welfare Act and the European Convention for the Protection of Vertebrate Animals used for Experimental and Other Scientific Purposes.

### Surgery and electrode implantation

All 10 rats were implanted with Neuropixels silicon probes targeting the MEC-parasubiculum region (3 of the rats were implanted bilaterally). Additionally, 5 of the rats with unilateral MEC-parasubiculum probes were implanted with a second probe targeting the contralateral anterodorsal thalamus (1 rat), contralateral presubiculum (2 rats) or hippocampus (2 rats; 1 contralateral, 1 ipsilateral). Finally, 1 rat was implanted with 4 probes targeting all areas (anterodorsal thalamus and hippocampus bilaterally, right presubiculum and left MEC-parasubiculum). Neuropixels 2.0 multi-shank probes (prototype and commercial) (*72*) were used in 8 of the rats, while phase 3A single-shank probes (*73*) were used in 2 rats.

Probes targeting MEC-parasubiculum were implanted 4.2-4.7 mm lateral to the midline and 0.0-0.3 mm anterior to the transverse sinus, at an angle of 18-25 deg in the sagittal plane, with the tip of the probe pointing in the anterior direction. These probes were lowered to a depth of 4.1-7.2 mm, often passing through postrhinal cortex before entering parasubiculum. Anterodorsal thalamus probes were positioned vertically at ML 1.4 mm lateral to the midline, AP 1.9-2.1 mm posterior to bregma, and lowered to a depth of 7.5-8 mm. Presubicular probes were inserted ML 1.65-3.15 mm lateral to the midline, AP 7.8-8.0 mm posterior to bregma, to a depth of DV 7.3-9.4 mm, either vertically or at an angle of 20 degrees in the coronal plane, with probe tips pointing laterally. Hippocampal probes were inserted ML 1.4-4.5 mm from the midline and AP 1.9-5.0 mm posterior bregma, to a depth of DV 5.5-7.1 mm, either vertically or at an angle of 10 degrees in the coronal plane, with the tip of the probe pointing laterally. One of the rats with probes in hippocampus had also received injections of adenoassociated virus (AAV8) carrying the fluorescent marker mCherry and the HM4Di DREADD receptor bilaterally in the hippocampus 4 weeks prior to probe implantation. The recording included in the present study was obtained without any administration of DREADD agonist. The probes’ ground and external reference pads were connected to a skull-screw above the cerebellum with a silver wire, and the implant was encapsulated in dental cement. The detailed procedures for chronic Neuropixels surgeries have been described elsewhere (*72*).

Postoperative analgesia (meloxicam and buprenorphine) was administered during the surgical recovery period. Recordings began when rats had recovered and resumed normal foraging behavior, at least 3 hours after surgery.

### Electrophysiological recordings and tracking

Instruments and procedures were similar to those described for Neuropixels recordings used in previous studies in the lab (*11*, *48*, *72–74*). Briefly, neural signals were amplified (gains of 500 for phase 3A and 80 for 2.0 probes), filtered (0.3-10 kHz for phase 3A and 0.005-10 kHz 2.0 probes) and digitized at 30 kHz by the probe’s on-board circuitry. Signals were multiplexed and transmitted to the recording system along a tether cable. Acquisition and probe configuration was controlled with SpikeGLX software (https://billkarsh.github.io/SpikeGLX/).

A motion capture system – based on retroreflective markers on the implant, OptiTrack Flex 13 cameras, and Motive recording software – was used to track head position and orientation in 3D. The 3D tracking coordinates were subsequently projected onto the horizontal plane for estimation of 2D position and head direction azimuth. For chasing experiments, 1-3 additional reflective markers were placed on the bait to track its location. In 2 chasing experiments where the reflective markers were not reliably registered by OptiTrack, we used DeepLabCut(*75*) to track the location of the bait in the overhead video frames. Back markers were attached in one animal. An additional camera (Basler acA2040-90umNIR) was used to capture overhead infra-red video at 50 Hz. Overhead video frames were aligned to OptiTrack tracking data with an affine transformation between corresponding points in the video and tracking data. Timestamps from each data stream were synchronized by using an Arduino microcontroller to generate randomized sequences of digital pulses, that were sent to the Neuropixels acquisition system as direct TTL input and to the OptiTrack system and video camera via infrared LEDs placed on the edge of the arena (*11*, *48*).

### Behavioral procedures

Recordings were obtained while animals freely foraged in an open field, chased a rapidly moving bait in the open field, walked backwards in a corridor or in the open field, ran back and forth for rewards on a linear track, or slept. Most recordings were performed within the first week after surgery (full range: 0-151 days post-operatively). All behavioral tasks for a given animal were performed in the same recording room, often consecutively on the same day. During pre-surgical training, some of the rats were food-restricted, maintaining their weight at a minimum of 90% of their free-feeding body weight. Food restriction was not used in any of the animals at the time of recording.

#### Open-field foraging task

Ten rats foraged for randomly scattered food crumbs (corn puffs or cookies) in a square open-field (OF) box with an arena size of 150 × 150 cm and 50-cm high walls. The floor and walls of the arena were black, apart from a white cue card (width of 41 cm, height of 50 cm) attached centrally to one of the walls. The arena was placed centrally in a large room (16 or 21 m^2^) with visual access to background cues. At the time of recordings, all rats were highly familiar with the environment and the task (having experienced 10–20 training sessions prior to surgery, lasting at least 20 min each). Recording sessions lasted 23-141 min. During recording, the rats alternated between active foraging and short periods of spontaneous alert immobility. Running and immobility periods were analyzed separately (see section on *alert immobility*).

#### Bait-chasing (pursuit) task

Seven rats with MEC-parasubiculum implants (5 of them with additional probes in hippocampus, presubiculum and/or anterodorsal thalamus) were recorded as they chased a rapidly moving bait in the same open-field arena as used for free foraging (150 × 150 cm, same visual cues and lighting). The bait consisted of a piece of corn puff attached to the end of a string together with 1-3 reflective markers. The other end of the string was connected to a thin black rod held by the experimenter. During the task, the bait was moved around inside the open field arena by the experimenter. The bait was moved in a rapid and erratic manner to encourage pursuit. Once the rats caught the bait, it was taken away from the rat, and a new bait was attached to the string. All rats were habituated to the task before surgery and recordings.

Recording sessions with chasing lasted 6-51 min, and contained several chasing bouts and reorienting events (see section of *chasing bouts and reorienting events*)

#### Backward-locomotion task

Five rats with MEC-parasubiculum implants (4 of them with additional probes in hippocampus, presubiculum and/or anterodorsal thalamus) were recorded while they walked backwards either in a narrow corridor (2 rats), in the open-field arena (2 rats), or both (1 rat).

Two different corridor-based setups were used to elicit backward locomotion. The first setup consisted of a single dead-ended corridor that was placed inside the open-field arena. This corridor had a length of 80 cm, width of 5 cm and transparent walls with a height of 50 cm. The narrow width prevented the rat from turning, making backward locomotion the only way to get out of the corridor. The second setup consisted of a plus-shaped arena with four dead-ended corridors with tapered openings connected at the center. The corridors had a length of 45 cm, width of 6 cm and opaque walls with a height of 40 cm. In both setups, food crumbs were scattered inside the corridors to encourage the rat to enter them. Once the rats reached the end of a corridor, they typically walked backwards to get back out. The corridors were too narrow for the rat to turn. The rats were habituated to the corridor tasks before neural recordings.

In the open-field version of the backward-locomotion task, the rat was placed in the standard open-field arena while the experimenter held a corn puff in front of it. Once the rat bit onto the corn puff, the experimenter applied gentle counter-pull, which often resulted in the rat dragging the bait backwards across the arena, typically over a period of 2-5 seconds.

#### Linear track task

Four rats with MEC-parasubiculum implants were recorded as they ran back and forth on a 2 m linear track, as previously described (*11*). The track was fitted with reward wells at both ends, from which liquid rewards (chocolate-flavoured oat milk) could be delivered. Once the rat consumed reward at one end of the track, the other reward well was filled. Before surgery, rats were trained to shuttle back and forth until they completed ∼40 laps per training session.

#### Natural sleep

Sleep recordings were obtained from 7 animals with MEC-parasubiculum implants (4 of them with additional probes in presubiculum and/or anterodorsal thalamus). Sleep was promoted by putting the rat in a black acrylic box (40 × 40-cm floor, 80 cm height), lined with towels on the floor, with room lights on, free access to water and pink noise played in the background to mask other sounds. The box walls were infrared-transparent, allowing the rat’s position and orientation to be tracked through the walls. Sleep sessions typically lasted 2–3 h.

### Spike sorting and single-unit selection

Spike sorting was carried out with KiloSort 2.5 (*72*), with customizations and unit selection procedures as previously described (*11*, *48*). Briefly, units were selected for further analysis on the basis of mean firing rates and their ‘waveform footprint’ (the spatial extent of spike waveforms), in order to exclude low-firing units, fast-firing interneurons and contaminated clusters. Standard inclusion criteria were mean firing rate of 0.1-10 Hz and waveform footprint <35 mm. Some thresholds were adjusted to account for region-specific firing properties. To account for the sparse firing of hippocampal cells, firing rate thresholds were set to 0.025 (lower bound) and 5 Hz (upper bound) for units in this region. To account for the high peak firing rates of anterodorsal thalamus head direction cells, firing rate thresholds were set to 0.1 and 60 Hz for units in this region. Additionally, hippocampal and thalamic units with waveform footprints up to 50 mm were included. Units that were recorded on sites located outside the regions of interest (MEC-parasubiculum, hippocampus, anterodorsal thalamus, presubiculum and postrhinal cortex) were excluded from further analysis, but units in transition zones were included.

### Preprocessing and temporal binning

Spike times were binned in 10-ms time bins for all population analyses, and tracking data was resampled at the same time intervals to align it with the spike-count data. For computational reasons, recording sessions were truncated in length to the nearest multiple of 100 seconds, by trimming the tail end of the behavior session.

### Rate maps and angular tuning curves

To generate 2D rate maps for the open field arena, position data were binned into a square grid of 2.5 × 2.5-cm bins. For each bin, we calculated each cell’s firing rate (number of spikes in the bin divided by time spent in the bin). Rate maps were smoothed with a cross-validated smoothing procedure, as previously described (*11*). The width, σ, of a gaussian smoothing kernel was optimized to minimize the mean squared error of the firing rate prediction (using the MATLAB function *fminbnd* and 1 *cm* < σ < 50 *cm*). Spatial autocorrelations and grid scores were calculated based on the individual cells’ rate maps, as described previously (*76*),

Angular tuning curves with respect to head direction, theta phase, internal direction (see section *Iterative decoding of internal direction based on population vector correlations*) or toroidal phase (see section on *single-module decoding*) were calculated by binning the angular variable into 60 evenly spaced angular bins. For each 6-degree bin, firing rate was calculated as the number of spikes divided by time spent in the bin. Angular tuning curves were smoothed with the same cross-validated smoothing procedure as the spatial rate maps (0.01 *rad* < σ < 1 *rad*), except for toroidal phase tuning curves, which were smoothed with a fixed-width gaussian kernel (σ = 5 bins).

### Identification of grid cells and grid modules

Grid cells and grid modules were detected using a clustering-based approach that identified groups of MEC-parasubiculum cells with similar spatially periodic activity, following a previously described procedure (*11*). Briefly, 2D autocorrelograms were computed from coarse-grained open-field rate maps (10 × 10 cm bins) and the central peak was masked. Vectorized autocorrelograms were concatenated in a matrix. Considering each autocorrelogram as a point in high-dimensional space, the Manhattan distances between all point-pairs were computed, each point’s 30 nearest neighbors were identified, and the resulting neighborhood graph was used as input to the Leiden clustering algorithm (resolution parameter of 1.0, or 1.5 for sessions with >1,000 units). Clusters of cells with clear and consistent grid patterns were then identified. For each cluster, we computed a median autocorrelogram across all of its member cells and used this to assess the cluster’s grid periodicity (by computing the grid score (*76*) of the median autocorrelogram) and its consistency (by correlating the autocorrelograms of each member cell with the median autocorrelogram, and taking the median across member cells). Clusters with a grid score >0.3, consistency >0.5 and ≥10 cells were classified as grid modules. Clusters with highly correlated autocorrelograms (r > 0.7) were merged.

### Classification of direction-tuned cells

Cells were classified as head direction (HD) cells if their HD-tuning curves differed significantly from a uniform distribution (p<0.001, Rayleigh test for non-uniformity) and had a Rayleigh mean vector length > 0.3. Cells were classified as internal direction cells if their tuning curves, with respect to decoded internal direction (see section *Iterative decoding of internal direction based on population vector correlations*), differed significantly from a uniform distribution (p<0.001, Rayleigh test for non-uniformity) and had a mean vector length > 0.3 that exceeded the HD mean vector length. Cells that passed both sets of criteria were classified as internal direction cells.

### Theta phase estimation and spectral analysis

Theta phase and time-varying power spectra were computed from the population spiking activity of all units (including fast-firing, putative interneurons) within the MEC-parasubiculum region. Spike times were binned into 10-ms bins and the resulting spike counts were summed across neurons and smoothed with a Gaussian kernel (*σ* = 20 ms for theta phase estimation and *σ* = 10 ms for spectral analysis).

For theta phase estimation, summed spike counts were upsampled by a factor of 10 using spline interpolation and peaks were detected to identify successive theta cycles. Phase values were assigned to peak times (0, 2π, 4π, …) and linearly interpolated between peaks to obtain a continuous phase estimate for each time point. The phase corresponding to minimal firing activity was defined as zero.

For spectral analysis, a multi-taper time-frequency spectrum was obtained from summed spike counts using the Chronux toolbox (http://chronux.org/, function mtspecgramc). Time-frequency spectra were computed over frequencies ranging from 1-15 Hz, using overlapping 2-s windows (step size of 0.5 s), a frequency bandwidth of 1 Hz and 3 tapers. Instantaneous theta power was quantified as the summed spectral power in the 6-12 Hz band at each time step, and theta frequency was defined as the frequency with maximum power within this band. Time periods where the peak spectral power was outside the theta-band were defined as non-theta states and were excluded from further analysis.

### Decoding sweeps based on population vector correlations

Position was decoded from MEC–parasubiculum or hippocampal population activity using a population-vector (PV) correlation method, as previously described (*11*). For each 10-ms time bin, we computed the Pearson correlation between the instantaneous population vector and session-averaged reference vectors derived from spatial rate maps in the open-field arena. The decoded position was taken as the bin with the highest correlation at each time step. Sweeps were detected as near-linear, outgoing sequences of decoded positions within each theta cycle, following a sequence-detection procedure described previously (*11*). Briefly, candidate sweeps were identified as the longest sequence of consecutive bins with <20 cm displacement and <90° directional change, truncated to maximize net displacement from start to end. Sweep direction and length were measured with reference to the lowpass-filtered decoded trajectory, which was computed by decoding position from spikes emitted in the first half of each theta cycle. A sweep vector, s, was defined as the vector between the lowpass-filtered decoded position at the beginning of the theta cycle and the distal-most point of the candidate sweep. Sweep direction and sweep length were defined as the direction and magnitude of the sweep vector. Sweeps were included for further analysis if the sequence contained at least 4 points and had a goodness-of-fit of r² > 0.5 with respect to the sweep vector.

For several visualizations, sweeps were transformed into head-centered coordinates by subtracting the lowpass-filtered trajectory and rotating by head direction. Session-averaged sweeps were computed by interpolating each sweep to 50 linearly spaced time points from the beginning to the end of the trajectory, and averaging across sweeps that followed left– or right-directed sweeps.

### Iterative decoding of internal direction based on population vector correlations

Internal direction was decoded from the activity of internal direction cells using the same PV-correlation approach but using angular tuning curves for internal direction instead of spatial position rate maps. To derive tuning curves for internal direction — an internal signal that is not directly observable but correlated with head direction — we used a two-step iterative decoding and tuning-curve estimation procedure (fig. S3B). In the first step, we decoded direction from all MEC-parasubiculum cells by correlating head-direction tuning curves and instantaneous PVs, and taking the circular mean of correlation values at time *t* as the decoded direction α:

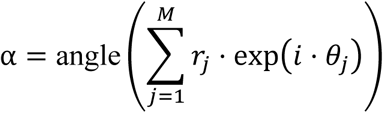

where θ_j_denotes the angular value of the *j*^th^ bin, *i* is the imaginary unit, *r*_j_ is the correlation value for the *j*^th^angular bin and *M* is the total number of angular bins. In the second step, we re-computed directional tuning curves with respect to the decoded signal. The resulting tuning curves were used to decode direction in the next iteration, before recomputing tuning curves once again. This procedure was repeated for three iterations. The tuning curves from the final iteration were used to decode internal direction from internal direction cells in downstream analyses. The time bin corresponding to the theta phase with maximal activity was used to express internal direction within each theta cycle. Although similar results were obtained using tuning curves with respect to tracked head direction for decoding (see *iteration 1* in fig. S3B), this iterative approach yielded sharper tuning curves and cleaner decoding (fig. S3B and S9C).

The same approach was used to decode direction from head direction cells in anterodorsal thalamus, presubiculum or postrhinal cortex.

For some analyses and visualizations (e.g. Fig 1f), we rotated the decoded direction (in allocentric coordinates) to a head-centered reference frame by subtracting the animal’s head direction.

### Computing the central axis of internal direction and detecting decoupling events

In this study, we decomposed internal direction dynamics into two components: (i) left–right alternation across successive theta cycles, quantified by an alternation width parameter, and (ii) a slowly drifting ‘central axis’, around which alternation was centered (Fig. 2A). The central axis of internal direction was estimated by smoothing internal direction across theta cycles with a gaussian kernel (σ = 1 theta cycle). Decoupling events – which were often observed during alert immobility – were defined as periods where the internal direction central axis deviated from head direction by more than 30° for more than 4 successive theta cycle. The central axis of sweeps, and decoupling of sweep direction from head direction, was computed in the same manner.

### Left-right alternation of sweeps and internal direction

The prevalence of left-right alternation of sweep direction or internal direction was quantified by identifying triplets of adjacent theta cycles where sweep direction or internal direction alternated in a left-right-left or right-left-right pattern (detected as sign-inversions in the angles between successive directions), and dividing the number of theta-cycle triplets with alternation by the total numbers of triplets where sweeps or internal direction were detected. The fraction of theta cycle triplets with directional alternation was compared to a shuffled distribution of alternation scores, where head-centered decoded directions were randomly shuffled (1000 iterations) before alternation in theta-cycle triplets were computed. Shuffled scores were typically in the range of 55-65%.

Alternation width (as reported in Fig. 1F, etc.) was quantified as the mean absolute angle between successive decoded directions, with analysis restricted to periods with left-right alternation.

Directional alternation was visualized in temporal autocorrelograms of angles between successive head-centered sweep direction or internal direction. Autocorrelograms were computed as the circular correlation between the original trace of head-centered directions and a series of lagged versions of the signal (lags from ±5 or ±7 theta cycles).

### Chasing bouts and reorienting events

The rats’ behavior in the object chasing task was segmented into chasing bouts and orienting events, based on the rats’ movement and orientation relative to the bait.

Chasing bouts were characterized by fast running towards the bait, and identified based on running speed and the offset between bait direction and head direction. Bait direction was defined as the direction of the vector from the rat’s head to the bait in allocentric coordinates (e.g. bait direction is 45° when the bait is located north-east of the animal’s head position). Chasing bouts were defined as periods where the rats were moving faster than 25 cm/s and the offset between bait direction and head direction was smaller than 30°. The number of detected bouts exceeding 0.5 s ranged from 29 to 279 per session, with durations up to 3.4 seconds.

During reorienting events, the rats lost track of the bait and paused, before making an orienting head movement towards the bait and starting a new pursuit towards. To identify these events, we first computed the absolute offset between bait direction and head direction, or the ‘head-centered bait angle’. Orienting events were identified as sudden reductions in head-centered bait angle, defined as negative peaks in its first derivative (<-2 rad/s), occurring as the rat transitioned from slow (<15 cm/s) to fast movement (>30 cm/s). Sweep direction and internal direction in the 350-ms time period leading up to orienting events were plotted against bait direction and head direction (e.g. Fig. 2C).

### Backward locomotion

Bouts of backward locomotion were identified as periods where the rats’ movement direction deviated from head direction by more than 2 radians (∼115°) for more than 500 ms with a movement speed >5 cm/s. Identified periods were curated by visual inspection of overhead videos.

### Alert immobility

Immobility periods with preserved theta oscillations during open-field foraging were detected as periods where the rat’s movement speed was lower than 3 cm/s and the peak of the time-frequency spectrum was within the theta-frequency range (6-12 Hz).

### REM sleep identification and classification

Periods of REM sleep were identified using a previously described sleep-scoring algorithm. Briefly, sleep was identified as continuous periods of immobility in the sleep box (longer than 120 s, locomotion speed below 1 cm/s, head angular speed below 6 deg/sec), and subclassified into REM sleep or slow-wave sleep based on the ratio of delta– and theta-rhythmic population activity in the recorded cells. 10-ms binarized spike counts from co-recorded MEC-parasubiculum cells were summed and band-pass filtered in the delta (1–4 Hz) and theta (5–10 Hz) frequency bands using a zero-phase, second-order Butterworth filter. Delta– and theta amplitude was calculated by Hilbert-transforming the filtered signal, taking its absolute value, and smoothing the resulting signal with a Gaussian kernel (σ = 5 s). REM sleep was classified as periods where the ratio of the z-scored theta– and delta amplitudes (theta/delta ratio) remained above 5.0 for at least 20 s were classified. REM sleep with slow– and fast theta oscillations were identified as periods where the instantaneous theta frequency was < 6.5 Hz and > 8.5 Hz, respectively. Theta surges during REM sleep were identified as peaks in instantaneous theta frequency that exceeded the 99^th^ percentile of theta frequencies across all REM periods in the session, with a minimum distance of 1 s between peaks.

### Decoding of direction and position signals during REM sleep

To decode direction signals from populations of direction-tuned cells in MEC-parasubiuculum, anterodorsal thalamus or presubiculum during REM sleep, we applied the same PV-decoding approach, using spike counts from REM sleep and tuning curves from a same-day open-field foraging session as reference.

To decode position from grid cells during REM sleep, we decoded position on the toroidal manifold of individual grid modules, using toroidal tuning curves from an open-field foraging session as a reference. Grid modules with >40 simultaneously recorded cells were included.

To construct toroidal tuning curves during open-field foraging, we first derived the first two grid axes **a**_l_ and **a**_2_ from the module’s median autocorrelogram, and used these axes as the basis of a toroidal coordinate system (fig. S8B). We next transformed the animal’s trajectory from Euclidian coordinates to toroidal phase angles, using the following formula:

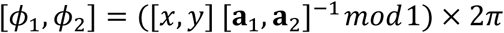

where [*x*, *y*] is the animal’s Euclidian position at time *t*, and [ϕ_l_, ϕ_2_] denotes the toroidal phase along grid axis **a**_l_ and **a**_2_, with values ranging from 0 to 2π. Angular tuning curves *f*(ϕ_l_) and *f*(ϕ_2_) with respect to the animal’s trajectory along grid axis **a**_l_and **a**_2_ were then computed as described in the section *Rate maps and angular tuning curves*.

We next decoded phase angles ϕ_l_and ϕ_2_in during REM sleep based on 10-ms population vectors and the tuning curves obtained above. Toroidal phases (ϕ_l_, ϕ_2_) were decoded independently using the Bayesian method, assuming Poisson firing and a flat prior:

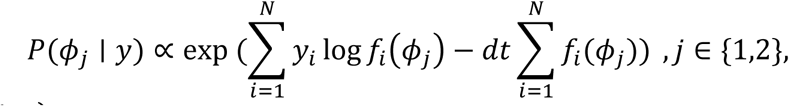

where *P*(ϕ_j ∣_ *y*) is the posterior for phase ϕ_j_, given spike counts *y* and tuning curves *f*(ϕ_j_). The decoded phase was taken as the bin maximizing *P*(ϕ_j_ ∣ *y*). Decoded phase angles were converted back to Euclidian coordinates on the grid unit tile, in the open-field reference space:

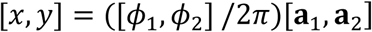

Decoding was carried out at slow and fast timescales to capture extended behavioral-timescale trajectories and sweeps, respectively. For slow decoding, spike counts were restricted to the first half of each theta cycle and smoothed with a Gaussian kernel of σ = 1.6 theta cycles. For visualizations (e.g. Fig. 5a), the slow decoded trajectory on the unit tile was unwrapped by hexagonally tiling the decoded position at each time step and selecting the point that minimized the distance to the previous unwrapped position coordinate. Single-module sweeps were detected in the fast decoded trajectory, as described for whole-population decoding (i.e. by finding consecutive time bins within each theta cycle where the decoded trajectory formed a smooth trajectory), except that spatial and directional offsets between consecutive decoded positions were computed with periodic boundary conditions.

### Histology and recording locations

The rats were euthanized with pentobarbital and perfused intracardially with saline followed by 4% formaldehyde. The extracted brains were stored in 4% formaldehyde, before being cut in 30-µm sagittal or coronal sections with a cryostat. The sections were Nissl-stained with cresyl violet and probe-shank traces were identified in photomicrographs. Recording sites on the probe were aligned to the histological sections by use of common reference points in histology and recordings (e.g., tip of the probe, intersection with brain surface or white matter, etc.). This alignment was used in conjunction with electrophysiological signatures (e.g. sharp-wave ripples, theta activity, axonal spikes in white matter tracts, etc.) and a stereotactic atlas (*77*) to determine recording locations. Visualizations of recorded brain areas were rendered with Urchin (https://github.com/VirtualBrainLab/Urchin).

### Data analysis and statistics

Data analyses were performed with custom-written scripts in Matlab and Python. Statistical analysis was performed in Matlab. Circular statistics were computed using the Circular Statistics Toolbox (*78*). Results are reported with means ± s.e.m. unless otherwise indicated. Statistical tests were nonparametric and two-tailed, unless otherwise indicated. Pearson correlations were used unless otherwise indicated. Power analysis was not used to determine sample sizes. The study did not involve any experimental subject groups; therefore, random allocation and experimenter blinding did not apply and were not performed.

**Fig. S1.**
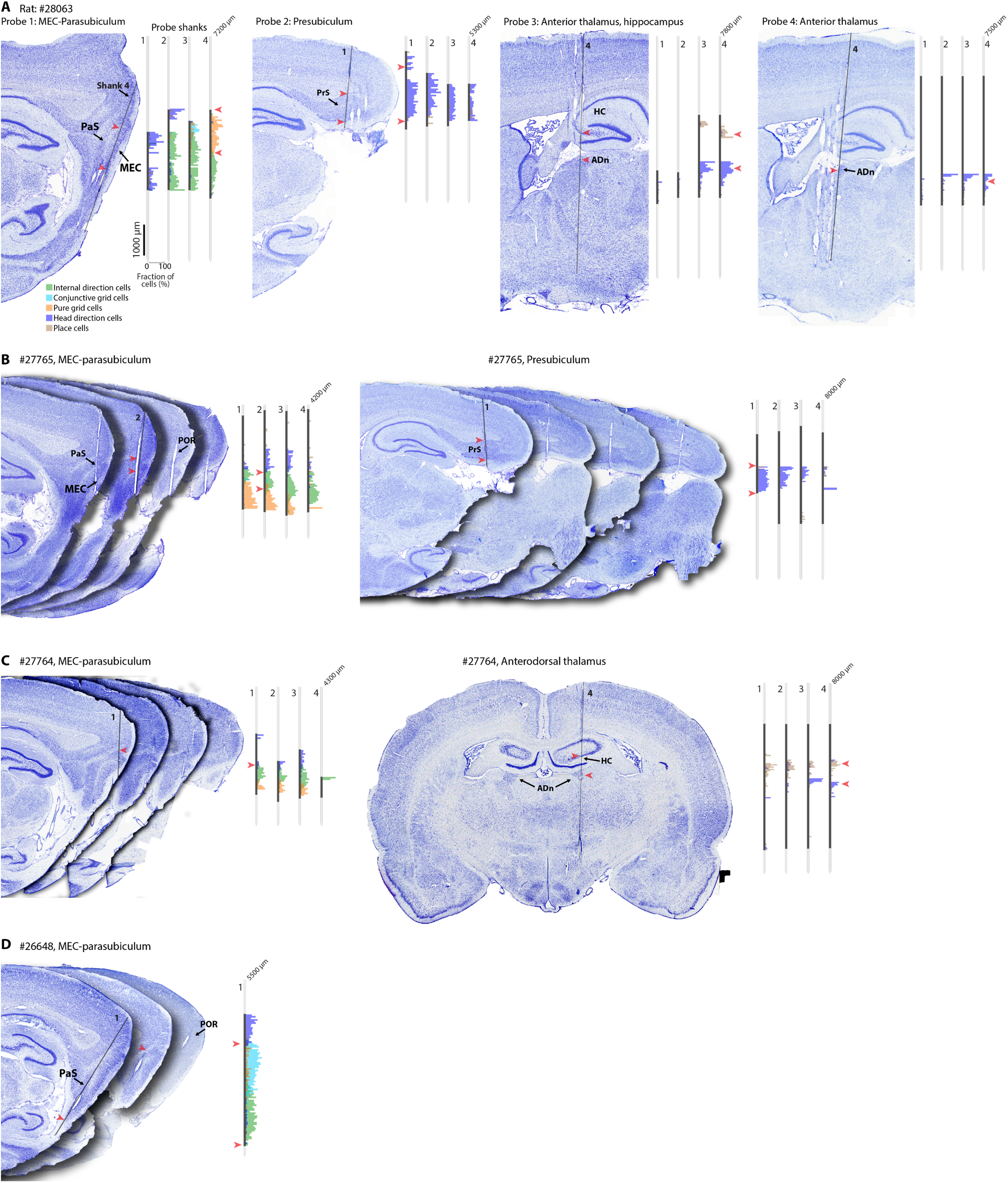
Histological sections and anatomical distribution of cell types. Panels show recording locations in four representative animals with at least two examples from each brain region included in the study. (**A**) Nissl-stained sagittal sections from a rat implanted with four four-shank Neuropixels 2.0 probes with probe tracks in left MEC-parasubiculum (MEC, PaS); right presubiculum (PrS); and bilaterally in anterodorsal thalamus (ADn) and hippocampus (HC). In each section, a black line indicates the trajectory of one of the shanks (indicated by number). Insets show fraction of functional cell types (cell type indicated by color) along each of the probe shanks (numbered), estimated by counting the number of functional cells recorded in 50 μm bins along the probe, divided by the total number of recorded cells in each bin. Cell counts are pooled across 10 recording sessions with different configurations of active recording sites (only 384 out of 5,120 sites could be recorded at any given time). Black portions of probe shanks show sites that were recorded from. Red arrowheads show corresponding locations in the histological sections and probe maps (probe 1, MEC; probe 2, PrS; probes 3 and 4, HC and ADn). (**B**) Histology and probe maps (as in (A)) for an animal with a in left MEC-parasubiculum and postrhinal cortex (POR; arrowheads at POR-PaS and PaS-MEC borders), and a second probe in right presubiculum. (**C**) Histology and probe maps for an animal with probes in left MEC-parasubiculum (arrowheads at the shank’s entry into parasubiculum) and right anterodorsal thalamus. (**D**) Histology and probe maps for an animal with a single-shank Neuropixels probe (same shank shown in multiple sections) in left MEC-parasubiculum, that also passed through postrhinal cortex (arrowheads at POR-PaS border and ventral PaS border).

**Fig. S2.**
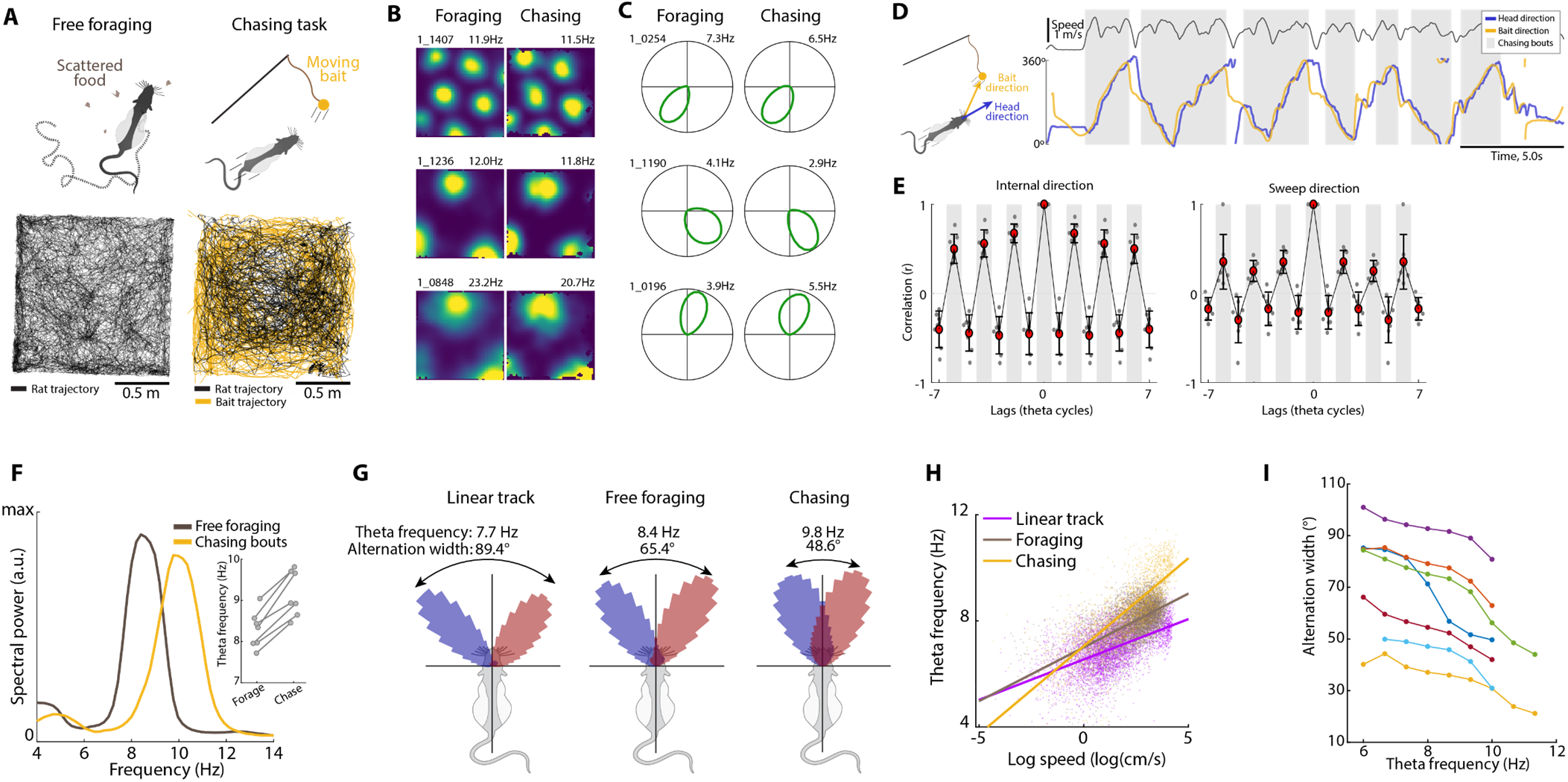
Sweeps and internal direction during chasing bouts. (**A**) Behavioral coverage during an example free foraging session (left) and a same-day chasing session in the same 1.5 x 1.5 m open field arena (right). The rat’s trajectory is plotted in black and the bait’s trajectory in yellow. (**B**) Spatial rate maps from three grid cells (rows) recorded during the free foraging session (left column) and chasing session (right column) shown in (A). Note that spatial tuning and firing rates are similar across conditions. (**C**) Internal direction tuning curves for three internal direction cells (rows) recorded during free foraging session (left column) and chasing session (right column) shown in (A). Note that directional tuning and firing rates are similar across conditions. (**D**) Left, schematic of the chasing task, with arrows indicating head direction (blue) and bait direction (the direction from the rat’s head to the bait; yellow). Right, tracked head direction and bait direction during a chasing session. Running speed is shown above. Note periods of rapid chasing directly towards bait (‘chasing bouts’; grey rectangles), interspersed with pauses (white gaps). (**E**) Temporal autocorrelogram of head-centered offsets in internal direction (left) and sweep direction (right) during chasing bouts (red dots, means; whiskers, s.d.). Note peaks at alternate theta cycles (indicated by grey and white background), consistent with prominent left-right alternation of the signals. Alternation was detected in 90.9 ± 1.9% of theta cycle triplets for internal direction (shuffled values of 66.2 ± 0.1%; mean ± s.e.m. across n = 7 rats), and 61.9 ± 1.9% of theta cycle triplets for sweep direction (shuffled values of 56.3 ± 1.2%). (**F**) Power spectrum of MEC-parasubiculum population spiking activity during free foraging in the standard open-field task (brown) and during bouts of chasing in the chasing task (yellow). Data is speed-filtered between 40 and 60 cm/s for both conditions. Note higher theta frequency during the chasing task (right-shifted peak in the theta-band). Inset, mean theta frequency during free foraging and chasing for all 7 animals (one line per animal). (**G**) Head-centered polar histograms of internal direction across tasks, for an example animal that was recorded while traversing a 2-m linear track, foraging in an open field, and chasing a moving bait. Red and blue histograms show the distribution of decoded directions when the previous decoded direction was oriented to the left (red) or right (blue). Mean theta frequency and alternation width are indicated above each plot. Note wide alternation and low theta frequency on linear track, narrow alternation and high theta frequency during chasing, and intermediate values during free foraging. (**H**) Scatter plot showing log running speed vs. theta frequency during linear-track running, open field foraging (‘foraging’) and chasing (same session as in (G)). Each data point corresponds to a 500-ms time bin. Lines of best fit are overlaid (linear regression). Note that theta frequency is higher during chasing compared to other conditions, even at comparable running speeds. (**I**) Average internal-direction alternation width, binned by theta frequency (bin width: 2 Hz), for all seven rats, each line corresponds to one rat. Note that alternation width decreases with increasing theta frequency (slope of –6.00 ± 0.89°/Hz, mean ± s.e.m.). Credit: rat (A, D and G), scidraw.io/Gil Costa.

**Fig. S3.**
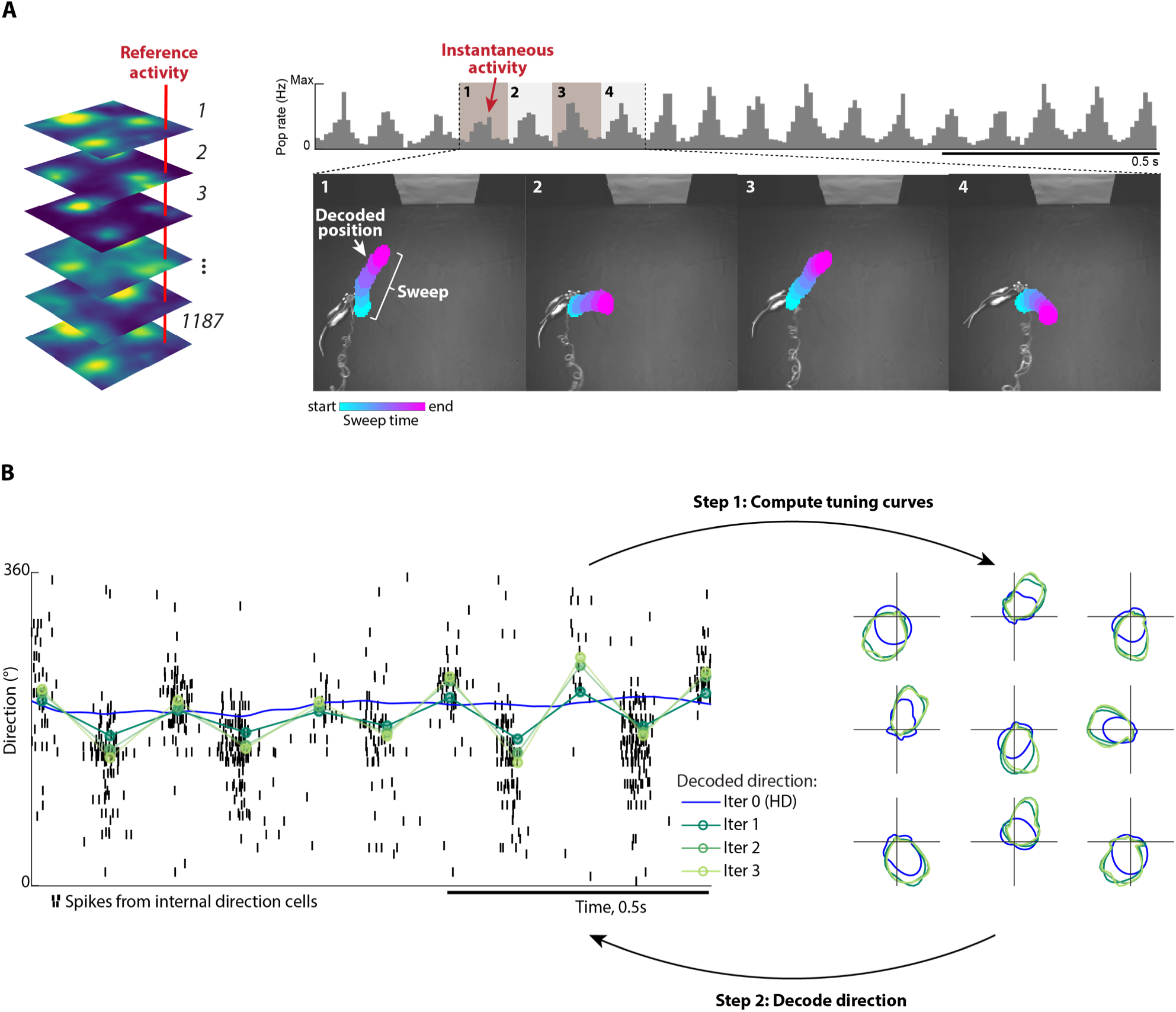
Extraction of sweeps and internal direction signals. (**A**) Methods for extracting sweeps from MEC-parasubiculum population activity. Left, stacked spatial rate maps from 6 example MEC-parasubiculum cells from an open-field foraging session with 1,187 co-recorded cells from this region. Each slice through the stack of rate maps (e.g. red line) forms a reference activity vector for a location in the arena. Top right, summed spike counts of all MEC-parasubiculum cells, with strong 8–10 Hz rhythmic population activity. Each bar corresponds to one 10-ms time bin. Bottom right, images show top-view snapshots of the recording arena at the beginning of four successive theta cycles. Decoded position, computed by correlating instantaneous population vectors and session-averaged population vectors for each location, is plotted as colored blobs. Position bins where correlation values exceed the 99.5^th^ percentile all bins are colored, with color indicating time within the theta cycle across 10-ms bins. Note that, within each theta cycle, the decoded position sweeps outwards from the animal in a left-right-alternating pattern. (**B**) Extraction of internal direction through iterative decoding and tuning-curve estimation. Left panel shows spikes from internal direction cells (black ticks), tracked head direction (blue) and decoded internal direction based on tuning curves from iterations 1-3 (green). Note that alternation becomes more prominent after the first iteration, providing a better fit to the spiking data. Right panels show representative directional tuning curves from 9 co-recorded cells, based on tracked head direction (blue) or decoded internal direction for iterations 1-3 (green). Note that tuning curves become sharper after the first iteration.

**Fig. S4.**
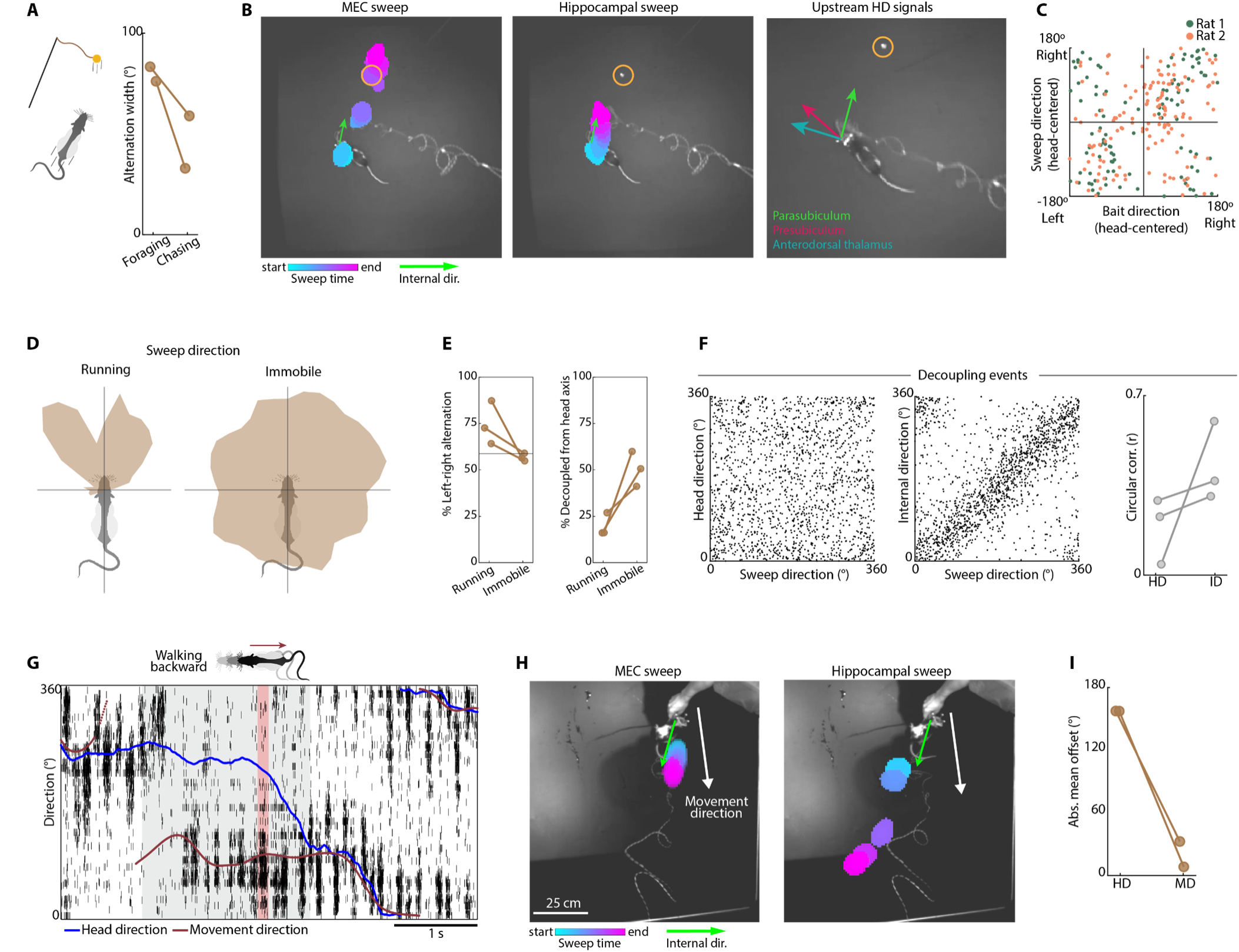
Hippocampal sweeps during chasing, backward walking and immobility. (**A-C**), Hippocampal sweeps during chasing. (**A**) Alternation width of hippocampal sweeps during free foraging and chasing bouts (n = 2 rats). Note narrowing of hippocampal sweep alternation in both rats (rat 1, from 76.1° to 32.9°; rat 2, from 83.3° to 58.9°). (**B**) Simultaneous decoding of position and direction across multiple regions of the navigation circuit during an orienting event in the chasing task. Left, decoded internal direction (green arrow) and sweeps (color blobs) from MEC-parasubiculum activity during a single theta cycle. Yellow circle indicates bait location. Middle, decoded sweep from hippocampal activity during the same theta cycle. Internal direction, decoded from parasubiculum cells, is shown in green. Right, decoded direction from parasubiculum (green), presubiculum (magenta) and anterodorsal thalamus (teal) during the same theta cycle. Note that the hippocampal sweep is directed towards the bait, like MEC-parasubiculum signals, while upstream head direction signals remain aligned to head direction. (**C**) Bait-directed hippocampal sweeps during MEC-parasubiculum decoupling events. Theta cycles where internal direction, decoded from MEC-parasubiculum activity, decoupled from the head axis were identified. Each dot shows hippocampal sweep direction and bait direction (both in head-centered coordinates) during one of these theta cycles. Data from 2 rats are shown in different colors. Note that hippocampal sweeps, like MEC-parasubiculum signals (Fig. 2), are directed towards the side of the bait in the majority of theta cycles (67.6% and 64.3% of theta cycles for rats 1 and 2). (**D-F**) Hippocampal sweeps during immobility with theta oscillations (related to Fig. 3). (**D**) Head-centered distribution of hippocampal sweep direction during running and immobility periods (with theta oscillations) in an example open field foraging session. Note that sweep direction is less rigidly aligned to the head axis during immobility, compared to running. (**E**) Left, fraction of theta-cycle triplets with left-right-alternation in hippocampal sweep direction during running and immobility (rat 1, 87.3% vs. 56.5%; rat 2, 72.6% vs. 59.0%; rat 3, 64.2% vs. 54.9% during running and immobility, respectively). Horizontal line at 50% indicates chance level for a random set of angles. Right, fraction of theta cycles where the internal direction axis decoupled by >30° from the head axis during running and immobility (rat 1, 16.2% vs. 60.0%; rat 2, 16.1% vs. 50.6%; rat 3, 26.9% vs. 41.1% during running and immobility, respectively). (**F**) Left panels, scatter plots of hippocampal sweep direction vs tracked head direction (left) or parasubicular internal direction (right), confined to immobility periods where internal direction is decoupled from head direction (data from one animal, one session). Note that hippocampal sweeps are aligned to parasubicular internal direction, and not head direction, during decoupling events. Right, circular correlation coefficients between sweep direction and head direction (r = 0.19 ± 0.07, p < 0.01 in 1/3 rats) or internal direction (r = 0.43 ± 0.09, p < 0.01 in 3/3 rats) for all 3 rats, one pair of points per rat. (**G-I**), Hippocampal sweeps during backward walking (related to Fig. 4). (**G**) Raster plot with spikes from internal direction cells during a period of backward walking (grey background). Note that population activity aligns with movement direction (brown) rather than head direction (blue) during this period. A single theta cycle is highlighted in red. (**H**) Mean offset between hippocampal sweep direction and either head direction or movement direction during all backward walking periods in 2 rats (rat 1, 157.3° and 8.2°; rat 2, 157.5° and 32.5° for head direction and movement direction, respectively). Credit: rat (A, D and G), scidraw.io/Gil Costa.

**Fig. S5.**
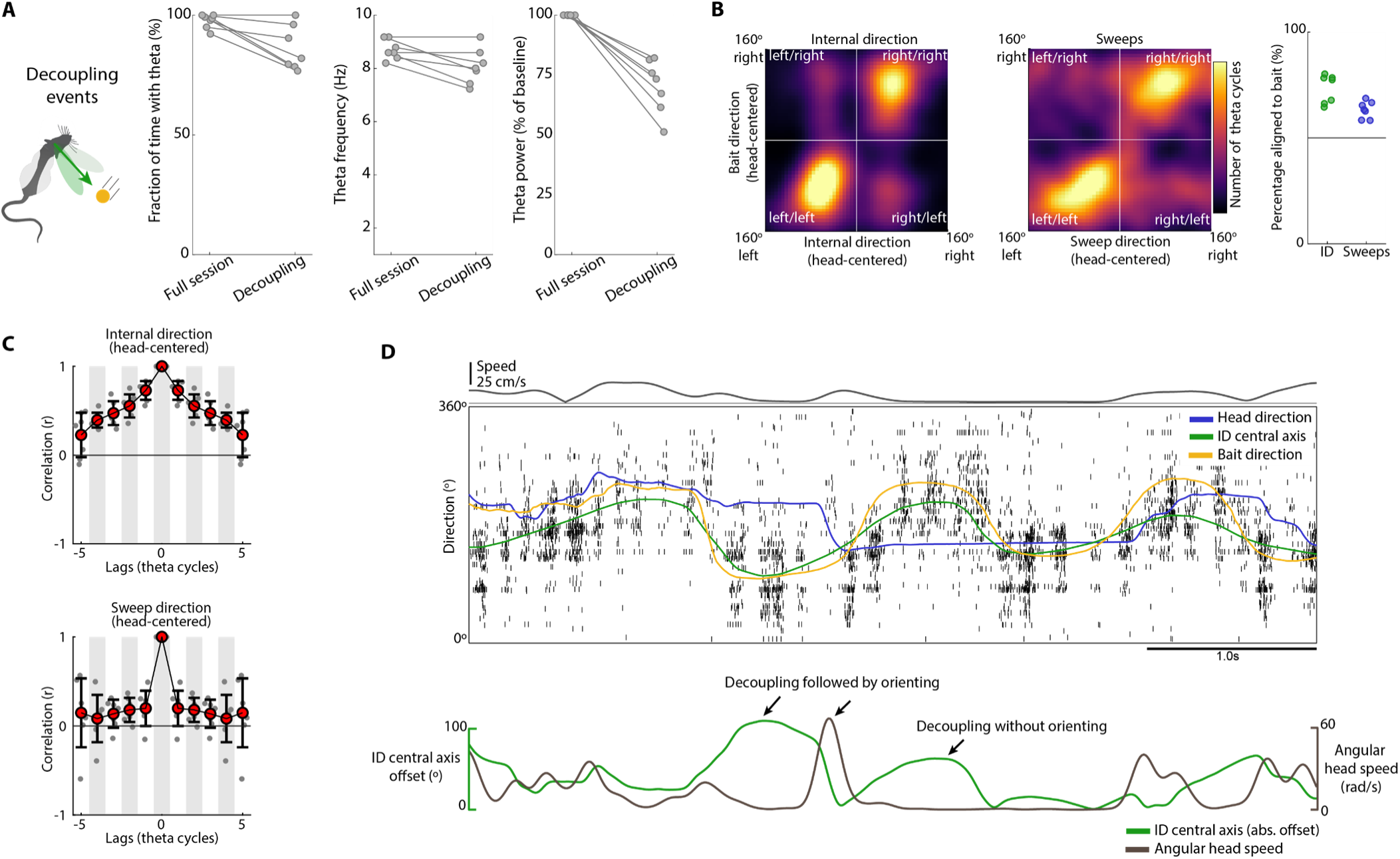
Theta oscillations, left-right alternation and behavior during decoupling events in the chasing task. (**A**) Fraction of recording time with theta oscillations (left), mean theta frequency (middle), and mean theta power (right) during the full chasing session and restricted to decoupling events (n = 7 rats, one pair of dots per rat). Note that theta oscillations are preserved during decoupling events (86.2 ± 3.5% of time in theta and theta frequency of 8.1 ± 0.3 Hz, mean ± s.e.m.). (**B**) Alignment of internal direction and bait direction (both with reference to rat’s head, as in Fig. 2A) during all ‘decoupling events’ pooled across all 7 animals (defined as periods where internal direction central axis deviates from the animal’s head axis by >30°). Note that internal direction (left panel) and sweep direction (middle panel) are directed towards the same side (right/left) as the bait during the majority of these events (73.2 ± 2.5% and 63.3 ± 1.5% of all decoupling events were ‘aligned’). Right panel shows alignment of internal direction (left) and sweep direction (right) to bait direction for all seven animals. (**C**) Temporal autocorrelograms of head-centered internal direction (left) and sweep direction (right) during decoupling events in the chasing task (red dots, means; whiskers, s.d.). Alternations are reduced or abolished with reference to chasing bouts (Extended Data Fig. 3e); they were observed in 61.0 ± 1.7% of theta cycle triplets for internal direction (shuffled values: 63.1 ± 1.7%), and 53.7 ± 1.2% of theta cycle triplets for sweeps (shuffled values: 46.1 ± 3.4%). (**D**) Top, raster plot with spikes from 697 internal direction cells during a segment of the chasing task where decoupling between internal direction and head axis is not followed by an orienting response (in most segments, decoupling is followed by immediate reorientation). Head direction (blue), bait direction (yellow) and internal direction central axis (‘ID central axis’; green) are also plotted. Running speed is shown on top. The animal follows a bait that swings back and forth in front of it. Bottom, angular head speed (brown) and offset between internal direction central axis and head direction (green) are plotted for the same segment below. Note that while internal direction tracks the bait throughout the segment, only one of the decoupling events is followed by an orienting response in this example.

**Fig. S6.**
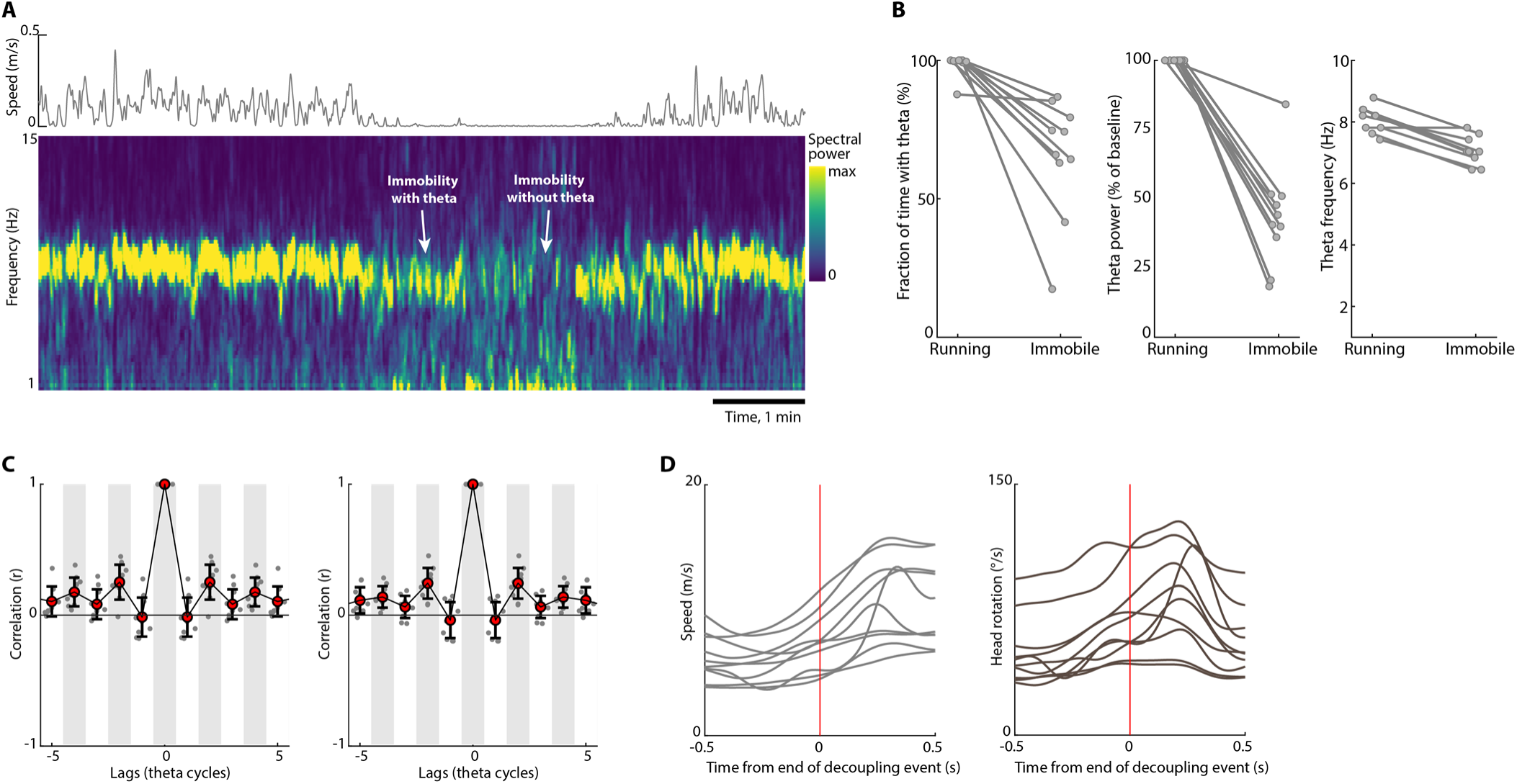
Internal direction signals during immobility anticipate upcoming behavior. (**A**) Example of immobility period with and without theta oscillations. Top, running speed during an 8 min segment of an open field foraging session with a long immobility period in the middle. Bottom, spectrogram of MEC-parasubiculum multiunit spiking activity during the same period. Note strong peak in theta-band during movement and a persisting peak in the theta-band during the first half of the immobility. (**B**) Fraction of recording time with theta oscillations (left), mean theta power (middle) and mean theta frequency (right) during open field running and immobility (n = 10 rats, one pair of points per rat). Theta oscillations were often present during immobility (65.3 ± 6.7% of immobility periods, mean ± s.e.m.), but with reduced frequency (7.1 ± 0.1 Hz) and reduced power (43.4 ± 5.8% of values during open field foraging). (**C**) Temporal autocorrelograms of internal direction in parasubiculum (left) and sweep direction in MEC (right) across successive theta cycles. Note absence of strong peaks at every other theta cycle, consistent with reduced left-right alternation during immobility. (**D**) Mean running speed (left) and angular head speed (right) triggered by end times of internal direction decoupling events (red vertical line) in all ten animals, one line per animal. Decoupling events were followed by a mean increase in running speed (6.7 ± 0.6 m/s vs. 10.3 ± 1.0 m/s before and after the end of decoupling, respectively, mean ± s.e.m.; increase of 53.3 ± 5.3%, Wilcoxon signed rank test: W(10) = 55, p = 0.002; latency to peak acceleration: 60.5 ± 32.3 ms). Decoupling events were also followed by increased angular head speed (53.3 ± 6.4 °/s vs. 76.2 ± 8.6 °/s, increase of 43.9 ± 6.1%, Wilcoxon signed rank test: W(7) = 55, p = 0.002; latency to peak angular head speed: 144.5 ± 23.4 ms).

**Fig. S7.**
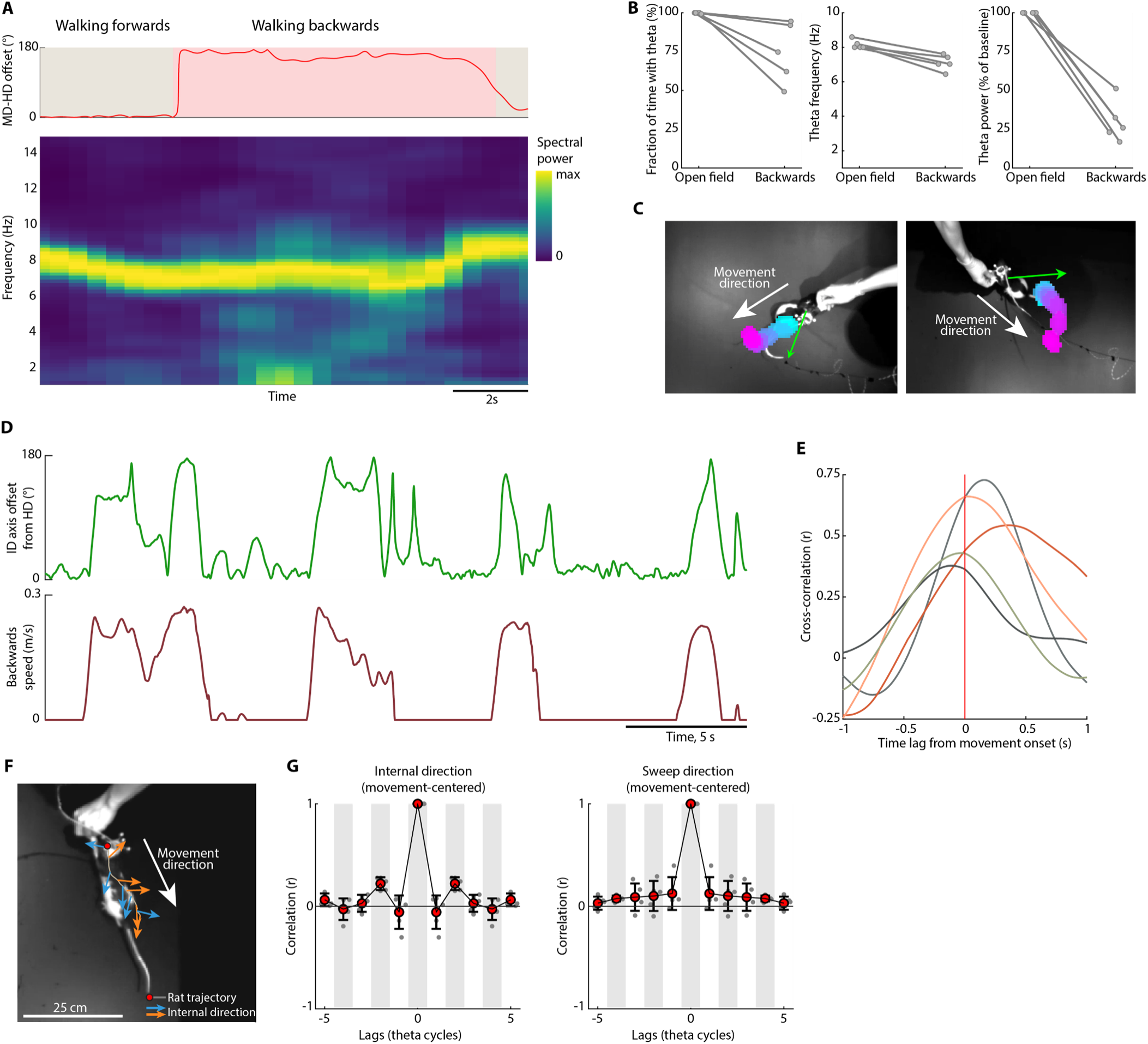
Central-axis inversion during backward walking is task-invariant and time-locked to movement. (**A**) Theta oscillations during backward locomotion. Top, offset between movement direction (MD) and head direction (HD) during a period of backward walking in the corridor (red shading). Bottom: Spectrogram of multi-unit activity during the same period (normalized by the peak density at each time point). Note preserved peak in theta range during backward walking. (**B**) Fraction of recording time with theta oscillations (left), mean theta frequency (middle) and mean theta power (right) during open field foraging (running periods) and backward walking (n = 6 rats, one pair of points per rat). Note that theta oscillations are present (74.5 ± 8.7% of backwards periods, mean ± s.e.m.), but at reduced frequency of 7.1 ± 0.2 Hz (compared to 8.1 ± 0.1 Hz in open field) and markedly reduced power (29.9 ± 5.9% of values during open field foraging). (**C**) Decoded internal direction and sweeps during two example theta cycle where the rat is dragging a piece of food backwards, against the experimenter’s pull. Note that sweeps and internal direction point backwards along movement direction, although the reward remains in front of the animal. (**D**) Inversion of the internal-direction axis during repeated bouts of backward walking. Top, offset between internal direction central axis and head axes during five bouts of backward walking. Bottom, backward movement speed during the same period. Backward speed was computed by projecting the animal’s velocity vector onto the backward head axis (180° from head direction) and rectifying the resulting signal. Note that the internal-direction axis inverts rapidly and persistently throughout each trial. (**E**) Cross-correlation between internal-direction axis offset and backward speed, computed over 2 s windows around axis inversions. Note cross-correlograms peaks near zero, indicating that backwards movement onsets coincide with axis inversions (non-significant lag of 8.0 ± 8.3 ms from movement onset to central axis inversion; Wilcoxon signed-rank test: W = 10, n = 5, p = 0.63). (**F**) Decoded internal direction during a period of backward locomotion where left-right-alternation is preserved. (**G**) Temporal autocorrelograms of movement-aligned internal direction (left) and sweep direction (right) across successive theta cycles during backward locomotion. Left-right alternation was observed in 63.2 ± 2.1% and 56.4 ± 1.7% (mean ± s.e.m.) of theta cycle triplets for internal direction and sweeps, respectively (shuffled values of 54.3 ± 2.1% and 51.3 ± 0.4%).

**Fig S8.**
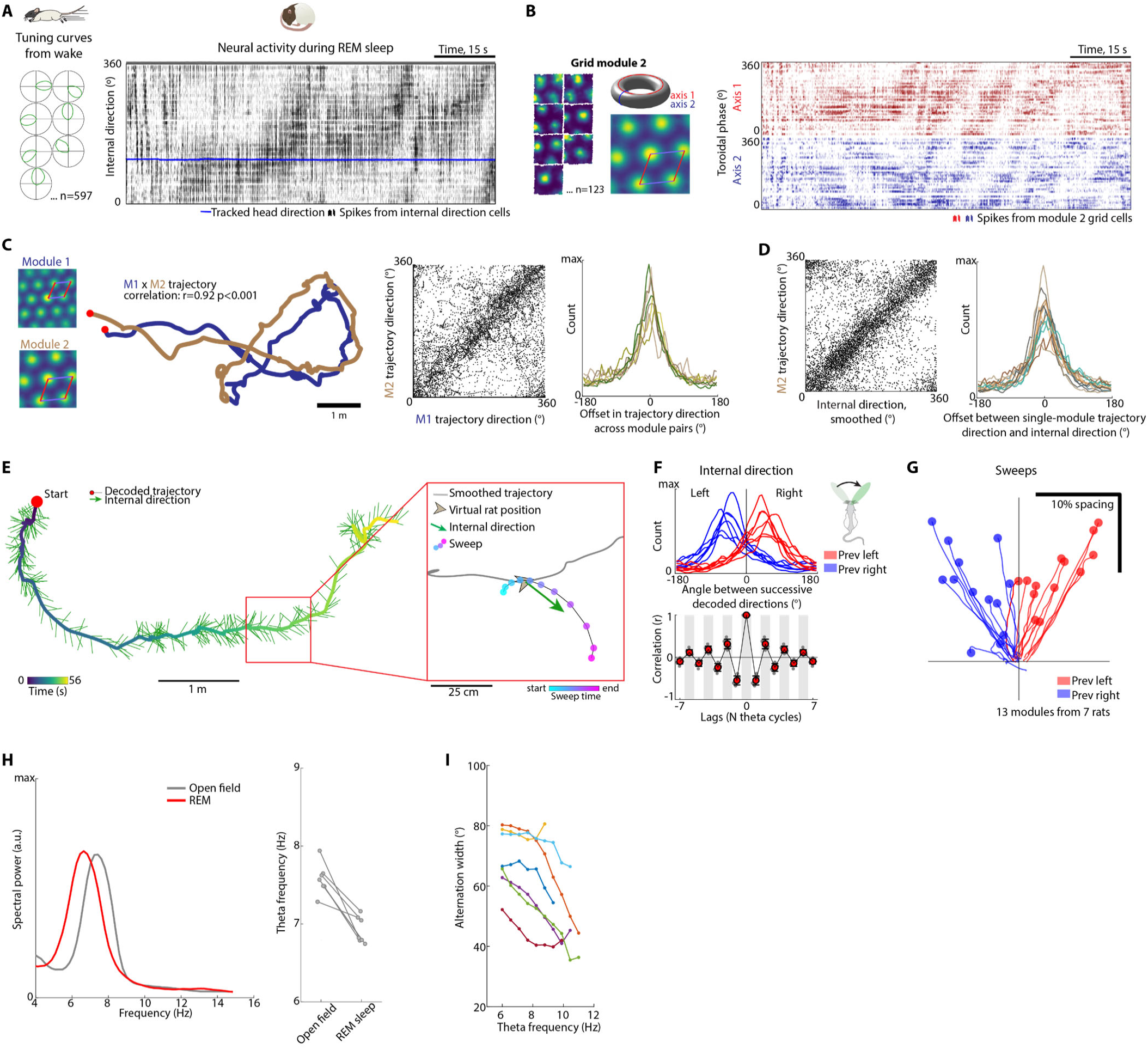
Population dynamics of internal direction cells and grid cells during REM sleep. (**A**) Left, example tuning curves for 7 out of 597 co-recorded internal direction cells during an open-field foraging session. Right, raster plot with spikes from all 597 internal direction cells during a subsequent 1-min episode of REM sleep. Cells are sorted along the y-axis by preferred direction during wake. Head direction is plotted in blue. Note wake-like dynamics of the internal direction signal over the course of the REM episode, with a packet of activity rolling around the ring-like manifold, despite absence of motor output. (**B**) Left, rate maps for 7 out of 123 grid cells from grid module 2 during the same open-field foraging session as (A). Median autocorrelogram of rate maps from all grid cells in the module is also shown. The first two axes of the grid pattern (red and blue lines) define a rhombus-shaped unit tile that is equivalent to a torus (inset). Right, spikes from all 123 grid cells in module 2 during the same REM sleep episode as (A), sorted by their preferred tuning along grid axis 1 (red) and 2 (blue). Note the slowly drifting packet of localized population activity, indicating that the population activity bump moves across the toroidal manifold throughout the REM sleep period. (**C**) REM sleep trajectories are aligned across grid modules. Left, spatial autocorrelograms and unwrapped decoded trajectories for modules 1 and 2 during the REM sleep period shown in Fig. 5A. Note that both grid modules play out the same extended 2D trajectory. Trajectory similarity is quantified by correlating the first derivative (movement vectors over 80 ms windows) of the trajectory from each module (mean ± s.e.m. trajectory similarity across all grid module pairs: r = 0.47 ± 0.03; p < 0.001 in 8/8 module pairs from 4 animals with >40 co-recorded grid cells in at least two modules). Middle, scatter plot showing movement direction on the toroidal manifold of modules 1 and 2 during all REM sleep episodes in one example animal (each dot corresponds to one 10ms time bin). Right, histograms showing alignment between trajectory direction during REM sleep in all module pairs (n = 8 module pairs from 4 rats). Each line corresponds to one module pair. Note that movement directions are aligned across modules (mean ± s.e.m angular offset of 3.49 ± 0.87°, n = 8 module pairs from 4 rats). (**D**) Left, scatter plot showing movement direction on the toroidal manifold of modules 2 and internal direction (smoothed across theta cycles) across all REM sleep episodes in one example animal (same session as (C)). Right, histograms showing alignment between trajectory direction and smoothed internal direction during REM sleep for all modules (n = 13 module pairs from 7 rats; each line corresponds to one module). Single-module trajectory direction is aligned with internal direction, with a mean ± s.e.m. offset of 6.9 ± 1.4°. Circular correlation between single-module trajectory direction and internal direction: r = 0.46 ± 0.03, mean ± s.e.m.; p < 0.001 in all 13 modules. (**E**) Left-right-alternating internal direction and sweeps are nested within extended REM sleep trajectories. Left, unwrapped trajectory decoded from grid cells of grid module 2 during a 56 s segment of REM sleep (same interval as in (A)), plotted as in Fig. 5A and color-coded by time. Decoded internal direction, from internal direction cells, are shown as arrows for individual theta cycles. Inset shows a sweep, decoded from grid cells of grid module 2, during a single theta cycle, with time indicated by color. (**F**) Top, distribution of angles between decoded direction at successive theta cycles (*n* = 7 rats, 1 session per rat) when the previous decoded direction was directed to the left (red) or right (blue). Bottom, autocorrelogram of decoded direction during REM sleep (red dots, means; whiskers, s.d.; shading indicates alternate theta cycles). Left-right-alternation of internal direction was present in 73.0 ± 2.0% of theta cycle triplets during REM sleep, mean ± s.e.m. across 7 rats. (**G**) Left-right-alternating sweeps in individual grid modules during REM sleep. Sweep trajectories from individual grid modules are referenced to the low-pass-filtered decoded trajectory, aligned to a ‘virtual head direction’ (low-pass-filtered internal direction) and averaged across theta cycles in which the preceding sweep went left (red) or right (blue). Data from 13 modules (with >40 co-recorded grid cells) from 7 rats are shown. (**H**) Left, power spectral density of MEC-parasubiculum multi-unit activity during open field foraging (grey) and REM sleep (red). Right, mean theta frequency during open field foraging (7.6 ± 0.1 Hz) and REM sleep across animals (6.9 ± 0.1 Hz). Theta frequency is lower during REM sleep compared to open field foraging (Wilcoxon signed-rank test: W = 0, n = 7, p = 0.016). (**I**) Internal-direction alternation width during REM sleep, binned by theta frequency (bin width: 2 Hz), for all seven rats, each line corresponds to one rat. Note that alternation width decreases with increasing theta frequency (slope of –4.0 ± 0.9°/Hz; correlation: r = –0.11 ± 0.03, p < 0.01 in 4/7 rats).

**Fig S9.**
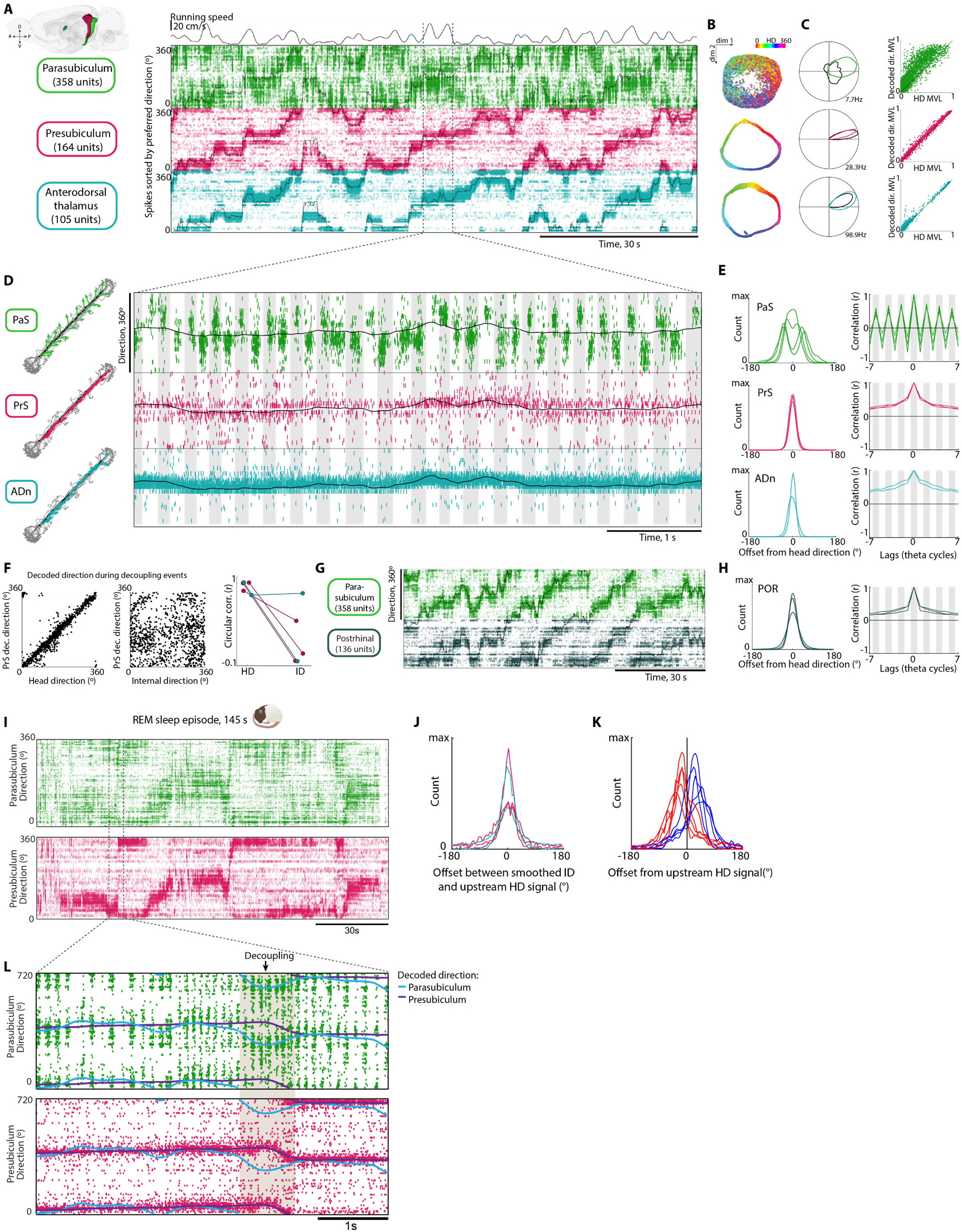
Population signals across nodes of the head-direction circuit during wake and sleep. (**A**) Raster plots showing spikes from direction-tuned cells in parasubiculum (PaS, green), presubiculum (PrS, magenta) and anterodorsal thalamus (ADn, teal) recorded simultaneously with four Neuropixels 2.0 probes in one animal during free foraging (ADn: 105 units, PrS: 164 units, PaS: 358 units; same animal as Fig. 6A, different session). Units are sorted by preferred direction and brain region (symbols as in Fig. 6A and B). Running speed is plotted at the top. Note that population activity in ADn and PrS is sharply centered around tracked head direction. **(B)** UMAP projection of population activity of directional cells in each region. Each dot corresponds to one time point, color-coded by tracked head direction. Note tighter coupling to head direction for the ring-shaped neural manifolds in ADn and PrS than in PaS. **(C)** Left column, tuning curves to tracked head direction (black) or decoded direction (green, magenta, teal) for an example cell in each region (top, PaS; middle, PrS; bottom, ADn). Note that the cell in PaS is more strongly tuned to decoded direction than tracked head direction, while tuning looks similar for the two signals in cells in ADn and PrS. Right, scatterplot of mean vector length (MVL) of tuning to tracked head direction (HD) or decoded direction for all cells in each region (n = 4 rats, 1 session per rat). Direction-tuned cells in parasubiculum are more strongly tuned to decoded direction than head direction (MVL 58.5 ± 2.6% larger than HD MVL, mean ± s.e.m., n = 1,387 cells), while mean vector lengths to head direction and decoded direction were similar for the two other regions (decoded direction MVL 5.3 ± 0.3% and 2.1 ± 0.4% larger than HD MVL, mean ± s.e.m. for PrS (n = 695 cells) and ADn (n = 153 cells), respectively). **(D)** Left, decoded direction from simultaneously recorded cells in PaS, PrS and ADn during traversal of a 2-m linear track. Note left-right alternation in the parasubicular internal direction signal, but not in upstream head direction signals. Right, raster plot as in (A), zoomed in on a 4-s segment where the rat is running straight. Tracked head direction is plotted in black. Note left-right alternation of the parasubicular signal, while signals in ADn and PrS remain stable. **(E)** Left, histograms showing offset between decoded direction and tracked head direction during open field foraging. Each line shows data from one animal (PaS, n = 4; PrS, n = 3; ADn, n = 2). Right, temporal autocorrelograms of head-centered decoding offsets for each region. Note precise head-direction representations and lack of left-right alternation in ADn and PrS. **(F)** Left panels, alignment between PrS-decoded direction and head direction (left) or internal direction (middle) during all immobility periods where the internal-direction centra axis was decoupled from the head axis. Data are from one example session; each data point corresponds to one theta cycle. Note that PrS-decoded direction remains aligned to head direction also in this condition. Right, circular correlation coefficients between PrS or ADn decoded direction and head direction or internal direction during decoupling events across all 4 rats. Each pair of connected dots corresponds to one cell sample from PrS (n = 3) or ADn (n = 2). Correlation between decoded direction and HD during decoupling events: r = 0.88 ± 0.03, mean ± s.e.m., p < 0.01 in 5/5 samples; correlation between decoded direction and ID during decoupling events: r = 0.25 ± 0.17, p < 0.01 in 2/5 samples. **(G)** Raster plots showing spikes from direction-tuned cells in PaS (green) and postrhinal cortex (POR, grey) during open field foraging. **(H)** Histograms of offset between decoded direction and tracked head direction and autocorrelograms of head-centered decoding offset, plotted as (E) with data from postrhinal cortex (n = 3). Note lack of left-right-alternation also in this region. **(I)** Raster plots showing spikes from direction-tuned cells in PaS (green; 221 cells) and PrS (magenta; 236 cells) during a 145-s REM sleep episode in an example sleep session. Note that internal direction is mostly centered around upstream head direction signals at this long timescale. **(J)** The internal-direction axis is aligned with upstream head direction signals during REM sleep. Histograms showing offset between decoded internal direction from PaS and decoded direction from upstream head direction areas. Both signals are smoothed across theta cycles with a gaussian kernel (s.d. = 1 theta cycle). Data from 4 rats are shown, each line corresponds to one paired recording from PaS and ADn (n = 2) or PrS (n = 3), mean ± s.e.m. offset of 1.08 ± 3.45° across all 5 samples. **(K)** Left-right-alternation of internal direction with respect to upstream head direction signals during REM sleep. Histograms show offset between internal direction (decoded from PaS) and upstream head direction signals (decoded from PrS or ADn) when the previous decoded internal direction was directed to the left (red) or right (blue) with respect to the upstream head direction signal. Data from 4 rats are shown, each pair of red and blue lines corresponds to one paired recording from parasubiculum and ADn (n = 2) or PrS (n = 3). **(L)** Internal direction transiently decouples from upstream head direction signals during REM sleep. Raster plots showing spikes from direction-tuned cells in parasubiculum and PrS during a 5-s segment extracted from (I) (same cells and sorting). Y-axis is repeated for visualization. Smoothed decoded direction from each region is plotted (PaS, light blue; PrS, purple). Note brief period where the parasubicular internal direction signal briefly decouples from the presubicular direction signal. These decoupling events occurred at a similar frequency during REM sleep (16.4 ± 5.2% of theta cycles, mean ± s.e.m. across 5 paired recordings from PaS and PrS or ADn) and non-speed-filtered open-field foraging sessions (18.1 ± 5.3% of theta cycles).

**Movie S1**.

Behavior and decoded internal direction during chasing of a moving bait. Part 1: Overhead video and 3D tracking during a representative segment of the chasing task, shown at real-time speed. Periods of rapid goal-directed running are interspersed with brief pauses during which the rat reorients towards the bait. Part 2: Decoded internal direction during a segment containing both pursuit and reorienting epochs (video slowed down). Internal direction is indicated by a green arrow, with arrow length proportional to spiking activity. Part 3: Same segment as in Part 2, but with internal direction smoothed across theta cycles to highlight the central axis of the internal direction signal (by averaging over left-right alternations). Note that internal direction follows the bait (orange circle) rather than the head axis before the onset of overt orienting movement. Part 4: Decoded sweeps (colored blobs) plotted along with decoded internal direction during the same reorienting event.

**Movie S2**.

Simultaneous decoding of population direction signals in parasubiculum (green) and presubiculum (magenta) during the chasing task. Decoded direction is smoothed with a 100-ms gaussian kernel to visualize the central axis of the alternating internal direction signal (by averaging over left-right alternations). Videos are slowed down. Part 1: Example where parasubicular internal direction decouples from the head axis, and the rat subsequently orients towards the bait. Parts 2 and 3 include examples of decoupling events without subsequent orienting movement. Note that the presubicular head direction signal remains aligned to the head axis, while parasubicular internal direction follows the swinging bait (orange circle).

**Movie S3**.

Internal direction and sweeps during immobility. Part 1: Decoded internal direction during a period with movement and immobility. Note food crumb that lands in the bottom right corner towards the end of the segment. Parts 2 and 3 show a single sweep decoded from MEC or hippocampus towards the food crumb in the corner. Part 4 shows sweeps decoded from each region side by side. Part 5: More examples of decoded internal direction (smoothed across theta cycles) events during immobility. Note that the internal direction central axis often points towards incoming food crumbs.

**Movie S4**.

Parts 1 and 2: Decoded internal direction (green arrow) and sweeps (colored blobs) as a rat walks backwards out of a narrow corridor. Video is slowed down. Part 3: Internal direction, smoothed across theta cycles and plotted with fixed arrow length, during several bouts of backwards walking. Video is played at real-time speed. Note that internal direction and sweeps point backwards, in opposition with the animal’s head direction.

**Movie S5**.

Simultaneous decoding of population direction signals in parasubiculum (green) and presubiculum (magenta) during backwards locomotion and passive movement. Decoded direction is smoothed with a 100-ms gaussian kernel to clearly visualize the central axis (while averaging over the left-right alternations). Part 1: Decoded direction during backwards walking in the open field. Part 2: Decoded direction during backwards walking in a narrow corridor. Part 3: Decoded direction during passive transport. Note that the internal direction signal in parasubiculum aligns with movement direction rather than head direction. The head direction signal in presubiculum remains aligned with head direction throughout the task.

